# On-tissue spatial proteomics integrating MALDI-MS imaging with shotgun proteomics reveals soy consumption-induced biomarkers in a fragile X syndrome mouse model

**DOI:** 10.1101/2021.11.09.467989

**Authors:** Min Ma, Qinying Yu, Daniel G. Delafield, Yusi Cui, Zihui Li, Wenxin Wu, Xudong Shi, Alejandra Gutierrez, Pamela R. Westmark, Meng Xu, Cara J. Westmark, Lingjun Li

**Affiliations:** School of Pharmacy, School of Medicine and Public Health, University of Wisconsin-Madison, Madison, Wisconsin, 53705, United States; Department of Chemistry, School of Medicine and Public Health, University of Wisconsin-Madison, Madison, Wisconsin, 53705, United States; Division of Otolaryngology, Department of Surgery, School of Medicine and Public Health, University of Wisconsin-Madison, Madison, Wisconsin, 53705, United States; Department of Neurology, School of Medicine and Public Health, University of Wisconsin-Madison, Madison, Wisconsin, 53705, United States; Molecular Environmental Toxicology Center, School of Medicine and Public Health, University of Wisconsin-Madison, Madison, Wisconsin, 53705, United States

## Abstract

Soy-based diets are associated with increased seizures and autism. Thus, there is an acute need for unbiased protein biomarker identification in Fragile X syndrome (FXS) in response to soy consumption. Herein, we present a spatial proteomics approach integrating mass spectrometry imaging (MSI) with label-free proteomics in a mouse model of FXS to map the spatial distribution and quantify the levels of proteins in the hippocampus and hypothalamus brain regions. In total, 1,004 unique peptides were spatially resolved, demonstrating the diverse array of peptidomes present in the tissue slices and the broad coverage of the strategy. A group of proteins that are known to be involved in the GABAergic system, synaptic transmission, and co-expression network analysis indicated that protein in soy group was significantly associated with metabolism and synapse modules in the *Fmr1*^*KO*^ brain. Ultimately, this spatial proteomics work laid the ground for identifying novel therapeutic targets and biomarkers for FXS.

## Introduction

Fragile X syndrome (FXS) is an X-linked neurodevelopmental disorder characterized by severe intellectual disability and other comorbidities, including autism, seizures, anxiety, and attention-deficit/hyperactivity disorder (ADHD)^1^. FXS is caused by the deficiency or absence of fragile X mental retardation protein (FMRP), an RNA binding protein (RBP) with a prominent role in the regulation of a large number of mRNAs in the brain and periphery ^2–4^. Given the critical role of FMRP in brain development and function, extensive studies have targeted the identification of FMRP binding partners, particularly at the transcriptome-wide level of the central nervous system (CNS)^2, 4–6^. Despite advances in understanding global translation and the identification of transcripts mis-regulated in the absence of FMRP, much remains to be learned regarding the proteins whose expression levels are altered, directly or indirectly, in FXS to improve our understanding of FMRP in brain neurodevelopment.

Our prior research demonstrates that single source, soy protein-based diets exacerbate seizures in mouse models of FXS, Alzheimer’s disease, and Down syndrome^7^, agreeing with numerous reports that detail the effects of diet on CNS function^8^. Soy-based diets have generally been considered a safe, healthy, and economical dietary alternative; however, these plant-based diets contain high levels of plant estrogens (phytoestrogens, isoflavones), which may mimic or antagonize natural estrogen activity and affect neuronal excitability. This may be particularly relevant to infant development if babies are fed soy-based infant formula, yet there is a dearth of studies regarding the effects of soy consumption on neurodevelopment^9^. Given the prevalent use of soy-based infant formulas, we conducted retrospective medical record and survey analyses, and found associations between the consumption of soy-based infant formula and increased incidence of seizures, autism, gastrointestinal problems, and allergies in children with autism and/or FXS^10–12^. The lack of molecular biomarkers correlated with the consumption of single source soy protein-based diets impede the understanding of how soy-based diets affect FXS.

Matrix-assisted laser desorption/ionization (MALDI) is a soft ionization technique used in mass spectrometry that allows the analysis of biomolecules that tend to be fragile and fragment when ionized by more conventional methods^13^. Mass spectrometry imaging (MSI) has become a powerful and successful tool in the context of detection and localization of a wide range of biomolecular species in recent years, including the imaging of lipids^14–17^, metabolites^18–19^, peptides^20–22^, proteins^23–26^, glycans^27–29^, neurotransmitters^30–31^, and drug compounds^32–33^. For protein MALDI-MSI studies, slices from fresh-frozen or paraffin-embedded formalin fixed tissue are mounted to microscope slides, followed by trypsin enzyme digestion to increase proteome coverage, aiding in protein identification^20, 34^. During a MALDI-MSI experiment, the laser beam moves across the surface of the matrix-covered tissue, which allows desorption and ionization of biomolecules^35^. However, for complex tryptic digestion peptides, it is difficult to confirm the identification of an individual peptide based solely on molecular mass. To solve this problem, some MALDI-MSI studies employ in situ tandem MS fragmentation analysis via tandem MS (MS/MS)^36–37^. However, the low fragmentation efficiency inherent to singly charged MALDI ions impairs identification. Liquid chromatography coupled with electrospray ionization (ESI) with tandem MS (LC-ESI MS/MS) offers front-end separation^38–39^ coupled with more efficient fragmentation capability due to multiply-charged ESI produced precursor ions, providing an attractive alternative to MALDI in situ MS/MS. Combining analyte localization with accurate mass measurements from MALDI-MSI with sequence identification by LC-MS/MS greatly enhances the breadth and depth of analysis of peptides from tissue digestion^40^.

Herein, we propose a comprehensive proteomics analytical workflow integrating MALDI-MS imaging with shotgun proteomics to discover and characterize biomarkers induced by high dietary consumption of soy protein (**Figure 1A**). We employed MALDI-MS imaging to analyze brain tissue from casein and soy protein-fed, wild type and *Fmr1*^*KO*^ mice, revealing thousands of spatially resolved precursor masses, which were putatively assigned to tryptic peptide identities. Candidate identities were then confirmed through extracting proteolytic peptides from adjacent tissue samples and subjecting them to LC-ESI MS/MS analysis. In total, we spatially resolved 1,004 unique protonated peptides from over 350 proteins, demonstrating the peptidomic diversity present in the tissue slices and the sensitivity of our technique. A shotgun proteomics strategy was then applied to extract protein candidates to quantify the relative abundance of different proteins revealed in MALDI MSI experiments. This strategy identified aberrantly expressed proteins caused by soy diet consumption that can be linked to the Alzheimer’s disease (AD) LOAD gene set, which illustrates the potential relationship between FXS and AD. In addition, co-expression network analysis showed that the *Fmr1*^*KO*^ genotype and soy protein-based diet were significantly associated with metabolism and synapse modules, via comparison of proteins in a mitochondrial module with glycolytic enzymes and the enzymes required for tricarboxylic acid (TCA) cycle. Finally, our annotated peptide sequence from the MALDI MSI workflow was linked to global shotgun proteomics illustrating the complementarity of multi-omics. To our knowledge, this work is the first systematic application of spatial MALDI-MSI proteomics, combining spatial mapping and shotgun proteomics, to identify soy-induced biomarkers, which have important implications for the discovery and identification of therapeutic targets and biomarkers for FXS.

**Figure 1.**
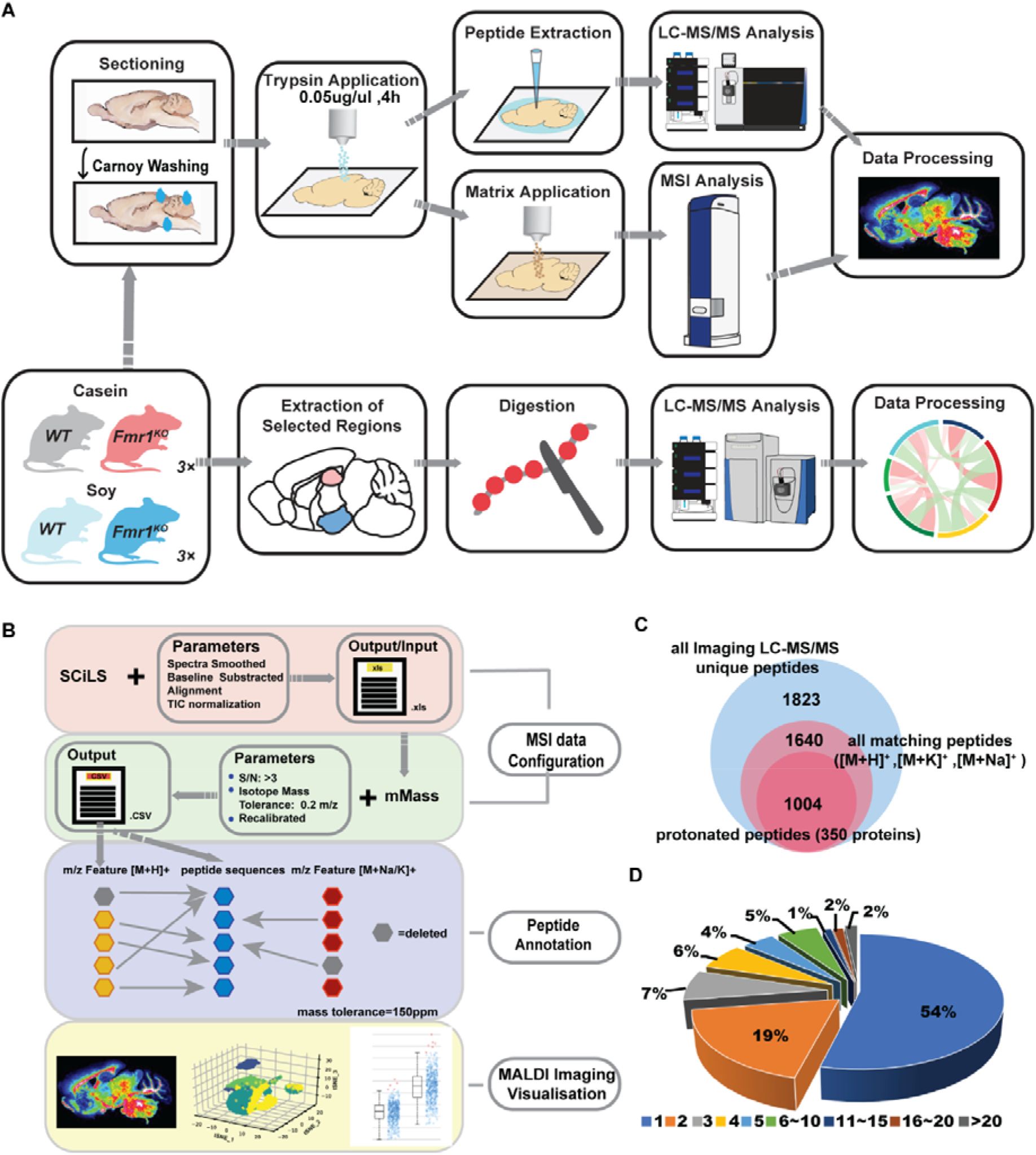
Multifaceted MS-based proteomic analysis of soy consumption in a FXS mouse model. **(A)** Schematic representation of the experimental strategy applied to resolve the spatial distribution of proteome (upper panel) and quantitative landscape of brain HPT and HPC proteome (lower panel) of soy consumption from WT and *Fmr1*^*KO*^ mice. **(B)** Graphical illustration of automated annotation of spatial proteomic mass spectrometry imaging data. The raw files utilize for peak picking in mMass software exported from SCiLS software. Parameters including imaging data pre-processing before exporting from SCiLS, and S/N, isotopic mass tolerance when finding peaks via mMass (upper two-panels); peptide annotation utilizes a peptide grouping strategy that deleted peptides which have larger mass errors or adduct peptides matched by one protonated peptide (lower middle panel); visualization of spatially clustered peptide ion images, representing the summary protein spatial distribution across whole mice brain tissues (lower panel). **(C)** The numbers of identified on-tissue LC-MS/MS unique peptides (class1); all matching peptides, including protonated and adduct peptides (class2); and unique matching protonated peptides (class 3) in MALDL-MSI. See supplementary Table S3 for more detailed information. **(D)** Pie chart depicting the percent distribution of multiple peptides assigned to one protein. The number of peptides for one protein are labeled with different colors from light blue (one peptide assigned to one protein) to dark gray (more than 20 peptides assigned to one protein).

## Results

### Construction of a robust mouse brain tryptic digestion imaging pipeline

Matrix-assisted laser desorption/ionization mass spectrometry imaging (MALDI–MSI) is a technique that enables label-free mapping of molecular species directly in tissue sections^41^. Although significant progress has been made in MALDI-MSI in recent years, the analysis of proteins is still challenging due to lower signal intensity, low confidence annotations, and biased localization within the image. The superposition of shifted masses from individual pixel peaks degraded the resolution and the mass accuracy in the average spectrum. Therefore, this approach is often unsuccessful in small regions of interest (ROIs) and for *m/z* peaks with low intensities.

An appropriate sample washing method is an important aspect of sample preparation to enhance target protein signals. For example, proper concentration of organic solvents is critical for eliminating salts, lipids, and metabolites that can reduce signals due to ion suppression^21^. Although there are many strategies for preparing imaging tissue sections^21^, it was necessary to design specific sample preparation methods to increase the signal of the targeted peptides in this study. In the workflow (**Figure 1A**, top panel), Carnoy’s solution (30% chloroform, 60% ethanol) was used to wash gelatin-embedded tissues. Keeping the same high humidity condition and the same concentration of trypsin enzyme (0.05 μg/μL), Carnoy’s solution produced a significantly higher signal to noise (S/N) ratio based on Wilcoxon signed-rank test (* *p*-value<0.01, ** *p*-value<0.001, ns-no significance) compared to the original washing method (70% ethanol, 100% ethanol) among three biological replicates (**Figure S1A**).

Tryptic peptides were extracted from two consecutive sections of each brain tissue and separated and analyzed by LC-MS/MS. Usually, numerous candidates were retrieved for a given mass of interest in LC-MS/MS. To achieve automated post-acquisition, an in-house script to match the MALDI-MSI peaks to the tissue LC-MS/MS results was used. The entire matching workflow is divided into three parts, as illustrated in **Figure 1B**. First, ROIs containing the entire area of the coronal section were selected in flexImaging. Then the raw files were imported to commercial software, Bruker’s SCiLS Lab, for spectra smoothing and alignment^42^, and total ion intensity (TIC) normalization. For the global pre-processing, the overall spectra for each of the analyzed sections obtained from SCiLS Lab were exported as comma-separated values (CSV) and read into the open-source software tool mMass. Mass spectral processing was performed using the following settings: (i) baseline subtraction, precision 25, relative offset 5; (ii) peak picking, S/N ≥ 3, relative intensity ≥ 0.5%; (iii) deisotoping, isotope mass tolerance 0.02 *m/z*, isotope intensity tolerance 50%; and (iv) recalibration based on the trypsin autolysis products, mass tolerance as 150 ppm. The tissue-specific peaks were exported to Excel (Microsoft) to compare with on-tissue digestion LC-MS/MS results. In tissue, the proportion of peptide signal from [Na]^+^ adducts may be higher relative to the [H]^+^ adducts, depending on the tissue preparation procedure and natural salt concentrations in different tissues and tissue regions^43^. However, since [H]^+^ adducts are still likely to be considerably more abundant than [Na]^+^ adducts, in this instance, priority was given to keeping the protonated peaks, if a peak could be matched to [Na]^+^ adducts and [K]^+^ adducts at the same time. For the case where multiply protonated peptides were matched, priority was given to peptides with the smallest mass error. Finally, the final peak list was imported into SCiLS Lab for imaging visualization.

### Global annotation peptide identification of MALDI-MSI

Study cohorts included wild type (WT) and *Fmr1*^*KO*^ (KO) littermate mice fed casein (C) or soy (S) protein-based diets. Our peak picking strategy (**Figure 1B**) was employed to obtain non-redundant peptide identifications within each tissue section. The peak list of each tissue section was corrected by subtracting all peaks detected in the gelatin tryptic digestion peptides (**Supplementary Table 1**). This correction removed all *m/z* features originating from the gelatin embedding material. The resulting peak list was averaged among three biological replicates of each cohort in **Figure S1B**. Approximately 1,000 specific peaks were detected per cohort. To further annotate MS imaging peaks, all on-tissue extracted unique proteolytic peptide sequences (**Supplementary Table 2**, sheet 1) were matched with each biological deisotope peaking list according to the pipeline in **Figure 1B**. The overlap of peptide signatures between groups (**Figure S1C**) demonstrated that at least 66% of the unique sequences in each cohort can be annotated in two biological replicates where only [H]^+^ adducts were considered within mass tolerances less than 150 ppm (**Figure S1D**). Nearly 90% of unique peptides from LC-MS/MS can be matched with the imaging peak list (**Figure 1C**, **Table S3**, sheet1). Of note, 1,004 protonated peptides (**Supplementary Table 3**, sheet 2), corresponding to 350 proteins, were assigned as MS^2^ unique sequences (**Figure 1C**). Among those proteins, 54% had one peptide sequence assigned. Also, nearly 50% of proteins include more than two tryptic peptides (**Figure 1D**), such as myelin basic protein (Mbp), neurofilament light polypeptide (Nefl), neurofilament medium polypeptide (Nefm), glyceraldehyde-3-phosphate dehydrogenase (Gapdh), and microtubule-associated protein tau (Mapt). For those proteins, we compared the peptide distribution within tissue sections (**Figure 2A**). The spatial distributions of the peptides were consistent with the parent proteins in the whole mouse brain, but distinct distributions were found in different brain regions (**Figure 2B**). The peptidyl-prolyl cis-trans isomerase Fkbp1a, which has potent neuroprotective properties and regulates glutamate-induced, dendritic protein synthesis^44^, had the same distribution in cortex across cohorts (**Figure 2B**, fifth row). These results are consistent with the MaxQB database^45^, which illustrates the distribution of Fkbp1a in human tissues (**Figure S2A**). On the other hand, collagen alpha-1(I) chain (Col1a1), which is a component of type 1 collagen and mutations cause Ehlers-Danlos syndrome, had higher expression in the cerebellum in all cohorts but was only enriched in the olfactory bulb region from the casein *Fmr1*^*KO*^ cohort (**Figure 2B**, row six). In addition, we also compared the 350 candidates to the curated database of mouse brain synaptic proteins (Genes to Cognition database, G2Cdb) ^46^. This identified an exceptionally strong overlap (45% of candidate proteins) with the Postsynaptic proteome (PSP), which contains 1,121 proteins (**Figure S2B**). We used the DAVID Bioinformatics database to identify the top pathways enriched in our targets (**Figure S2C**; **Supplementary Table 3**, sheet3). Strikingly, the most significant overlaps included biosynthesis of antibiotics/amino acids, glycolysis, synaptic, PI3L-Akt signaling pathway, and GABAergic synapse, etc., which suggests a direct role of diet - in - regulating the FXS proteome, as well as cognitive and behavioral deficits. Finally, the tissue was successfully classified by performing non-supervised spatial segmentation analysis (**Figure S2D**). Taken together, this robust strategy not only has high reproducibility but also enables the exploration of distinct spatial distributions of protein markers. This information, combined with the annotated peptide sequences in the mouse brain, provides a powerful tool to identify potential biomarkers induced by soy protein consumption in *Fmr1*^*KO*^ mice.

**Figure 2.**
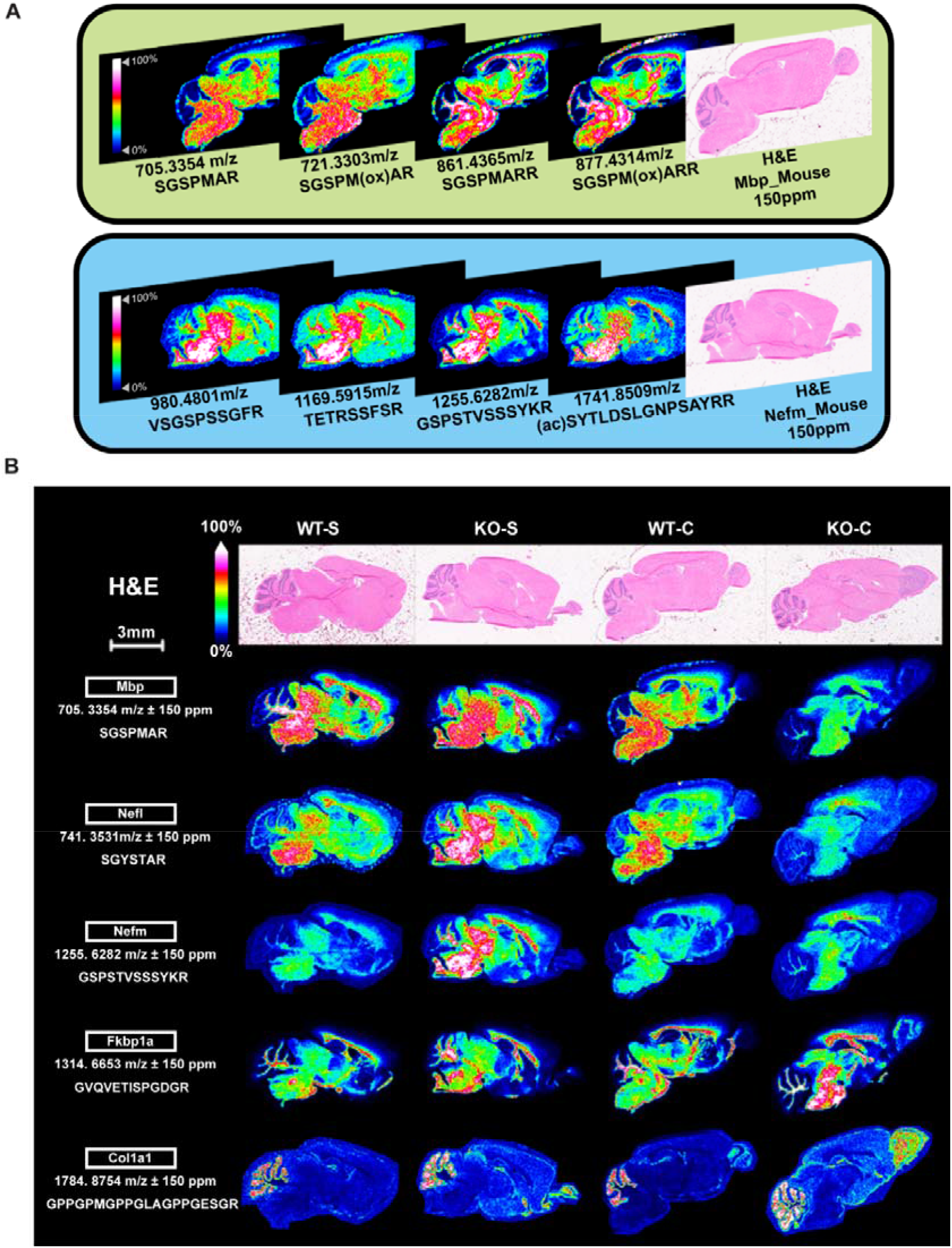
Overview of spatial distributions of the on-tissue digested peptides in positive ion mode. **(A)** Ion distributions representing Mbp peptides (upper panel) and Nefm peptides (lower panel) obtained from Bruker rapifleX MALDI-TOF/TOF were assigned to peptides identified by on-tissue tryptic digestion LC-MS/MS. Images were generated using SCiLS Lab. Hematoxylin and Eosin (H&E) staining images were shown on the right of the corresponding tissue section. **(B)** Summed ion spatial distribution of different representing protonated peptides. Images were generated using SCiLS Lab. H&E staining images were shown on top of the corresponding tissue sections.

### Spatial proteomics reveals candidate biomarkers in the hypothalamus and hippocampus

Feeding behavior is complex and modulated by various contextual factors and previous experiences^47–48^. The hypothalamus (HPT) plays key roles in appetite, food intake, whole-body energy homeostasis, glucose metabolism, and body weight regulation^49^. The hippocampus (HPC) controls fundamental learning and memory processes and is heavily influenced by feeding behavior^47^. To identify altered expression of protein biomarkers in HPT and HPC in response to the *Fmr1* genotype, the four treatment cohorts were integrated into two groups and analyzed separately, i.e., WT group (WT-C and WT-S) and *Fmr1*^*KO*^ group (KO-C and KO-S). To further narrow down the list of promising targets, we screened all-matching peaks (**Supplementary Table 3**, sheet1) to ensure that these peaks were detected in at least two biological replicates in all cohorts. Among these reproducibly detected peaks, we identified 350 unique protonated peptides from 178 proteins in WT group (**Supplementary Table 2**, sheet 2) and 380 unique protonated peptides from 196 proteins in *Fmr1*^*KO*^ group (**Supplementary Table 2**, sheet3). Comparison of the two groups of peptides and proteins revealed a high degree of overlap (73% proteins in all groups, **Figure 3A**). However, only 17.5% of proteins were specific in the *Fmr1*^*KO*^ group, including several well-known FMRP candidate genes such Prr12, Cplx2, Tubb3, etc. (**Supplementary Table 2**, sheet4).

**Figure 3.**
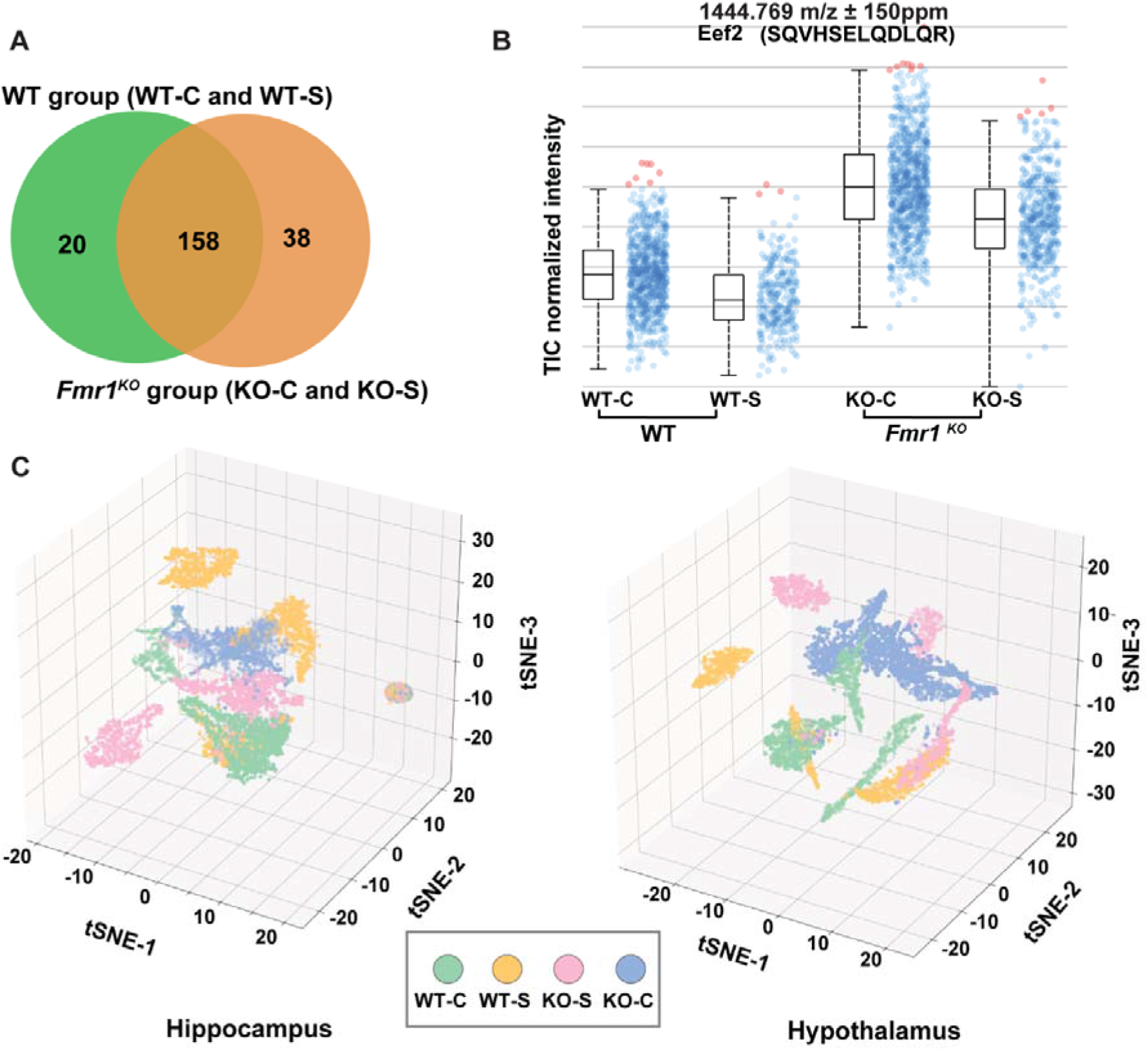
Annotated proteomes of FXS vs. WT mouse brain tissues. **(A)** Overlap in genotype groups between *Fmr1*^*KO*^ and WT when comparing two groups’ proteins among reproducibly detected peaks from at least two biological replicates from all four cohorts. **(B)** Normalized intensity of total ion signals of the Eef2 protonated peptide in four cohorts in HPC. The boxplot points correspond to relative intensities of *m/z* 1444.769-related peaks taken from each spectrum in all images acquired. **(C)** The t-distributed stochastic neighbor embedding (tSNE) visualization shows unsupervised annotated peaks clustering, revealing four distinct cohorts in HPC (left panel) and HPT (right panel).

FXS is the leading cause of autism spectrum disorders (ASDs), and 90% of affected males show some form of atypical behavior characteristic of autism^50^. Comparison of the protein-coding genes from the Darnell dataset^51^ and 213 candidate genes associated with ASDs from the SFARI autism database (https://sfari.org/resources/sfari-gene)^52^ with our dataset revealed varying degrees of overlap. A high degree of overlap was shown between our data and the Darnell database—29 (16%) and 36 (18%) targets in WT group and *Fmr1*^*KO*^ group, respectively (**Figure S3A**; **Supplementary Table 2**, sheet5 and sheet6), but only 4 and 6 targets were common between our dataset and the SFARI database for WT group and *Fmr1*^*KO*^ group, respectively (**Figure S3A**, **Supplementary Table 2**, sheet5 and sheet6). Among the targets in both datasets, eukaryotic elongation factor 2 (Eef2) is necessary for metabotropic glutamate receptor-mediated long-term depression (mGluR-LTD), a form of synaptic plasticity that requires de novo protein synthesis^53^. However, our imaging signal intensity showed that the level of Eef2 expression was moderately downregulated in the KO-S cohort compared to the KO-C cohort (p-value<0.01), even though Eef2 was elevated in *Fmr1*^*KO*^ group in comparison to WT group in HPC (**Figure 3B**). The ROC plot showed that Eef2 can be used to discriminate treatment cohorts (**Figure S3B** of WT group, **Figure S3C** of *Fmr1*^*KO*^ group). The same trend was also found in the HPT (**Figure S3D**). Thus, consumption of the single source soy protein-based diet is associated with decreased expression of Eef2 in both HPT and HPC in *Fmr1*^*KO*^ mice (compared to WT baseline), albeit the effect is moderate. The mechanism and biological significance of dysregulated Eef2 levels in FXS remain to be determined.

Given the complexity of comparing pixel-level intensity and distribution of assigned *m/z* values across tissue sections, we employed t-distributed stochastic neighbor embedding (t-SNE) to help inform the underlying similarity and differences between treatment groups. Full details can be found in the **Supplemental Information**. Briefly, using stained brain sections as reference, HPT and HPC brain regions were outlined in SCiLS Lab, the pixels lying within each region were normalized according to tissue-level total ion count and then exported for analysis. All HPT and HPC regions were concatenated into a two-dimensional array and coded by treatment group (**Figure S4**). The pixel-level intensity of each *m/z* value putatively assigned to a tryptic peptide was extracted and used to compile a three-dimensional training data set. After iterative optimization to determine appropriate t-SNE parameters, data points were clustered in three-dimensional space (**Figure 3C**). Slices from the same genotype cluster together more closely. The acquired peak list was imported in SCiLS Lab, and Student’s t-test analysis was performed. The Student’s t-test results showed that almost all peaks are significantly different (*p*-value<0.05) as a function of the *Fmr1* genotype (see **Supplementary Table 2**, sheet2 and sheet3).

Finally, we compiled a list of common targets in the two published databases and our imaging dataset, which represent a core set of synaptic proteins encoded by autism-associated genes, bound at the mRNA level by FMRP aberrantly expressed in response to soy-based diet (**Supplementary Table 2**, sheet 6). These proteins include Atp1a3, synapsin 1 (Syn1), and Slc1a2. Of note, the overlap among these datasets identifies biomarkers responsive to *Fmr1* genotype and soy consumption and thus potentially involved in the development of autistic phenotypes.

### Global shotgun proteomics analysis

MALDI imaging mass spectrometry serves as a useful tool that enables simultaneous detection and spatial mapping the distribution of putative peptide and protein signals; however, peptide identification is still a major challenge due to low S/N ratios, low fragmentation efficiency for MALDI-produced precursor ions, and limited quantification capacity between different experimental cohorts. To address these issues, we conducted parallel analyses on tryptic digests of homogenized mouse HPT and HPC by LC−ESI MS/MS. Samples were collected from four cohorts (WT-C, WT-S, KO-C, and KO-S) with 3 biological replicates each. Proteins were subjected to standard bottom-up proteomics workflows that involved tryptic digestion and label-free quantification to explore global protein abundance alterations and further correlate the findings with MALDI MS imaging results (**Figure 1A**). A total of 3,521 and 3,781 proteins (False Discovery Rate, FDR < 1%) were identified across the 4 cohorts in HPT and HPC, respectively (**Supplementary Table 4**, sheet 1 and **Supplementary Table 5**, sheet 1). Proteins were subjected to a binary comparison between soy protein versus casein protein diet in WT group and *Fmr1*^*KO*^ group. Those identified in at least two biological replicates in each cohort were considered quantifiable and resulted in 1866 (WT group; **Supplementary Table 4**, sheet 2), 2108 (*Fmr1*^*KO*^ group; **Supplementary Table 4**, sheet 3) quantified proteins in HPT and 2217 (WT group; **Supplementary Table 5**, sheet 2), 2061 (*Fmr1*^*KO*^ group; **Supplementary Table 5**, sheet 3) quantified in HPC (FDR < 1%, **Figure 4A**). We also examined the reproducibility of protein abundance across technical replicates (n=3) and biological replicates (n=3) using Perseus software^54^ (**Figure S5A** for HPT, **S5B** for HPC). Overall, pairwise comparisons of all technical replicates showed excellent quantitative reproducibility with a median Pearson correlation of 0.97 (**Figure S5**). A lower median correlation (*r* = 0.95) between biological replicates was observed, indicating a larger variation in biological replicates than in technical replicates.

**Figure 4.**
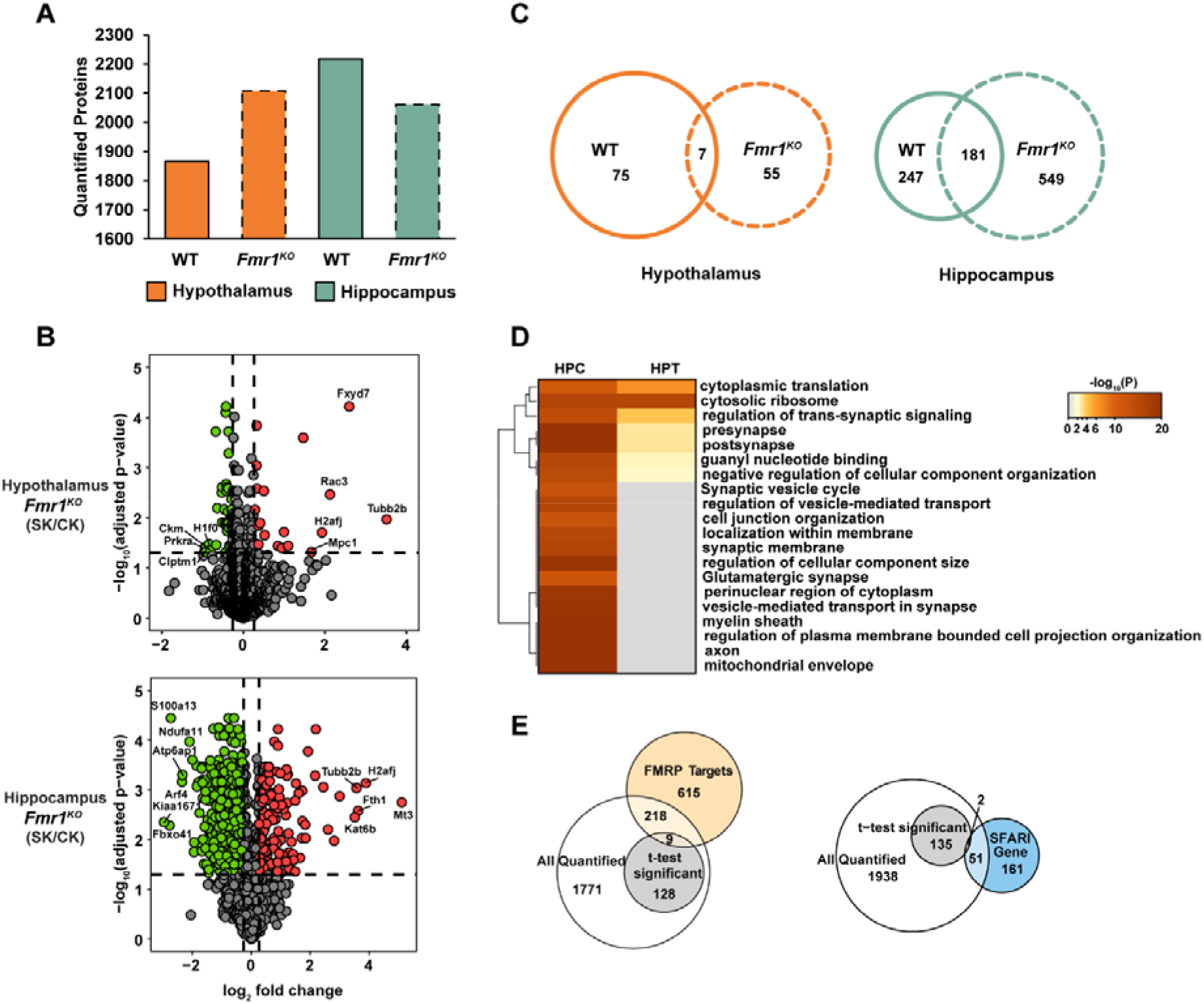
Global shotgun quantitative proteomic analyses of mouse HPT and HPC. **(A)** Number of proteins quantified using shotgun LC-MS/MS strategies in WT group and *Fmr1*^*KO*^ group in HPT and HPC, respectively. Bars with solid line border represent WT group and bars with dashed line border represent *Fmr1*^*KO*^ group. **(B)** Volcano plots showed protein expression level changes between soy protein versus casein protein diets in *Fmr1*^*KO*^ group for HPT (upper panel) and HPC (lower panel). Log_2_ protein fold changes are plotted against the negative log_10_ adjusted *p*-values. Points above the non-axial horizontal line represent significantly altered proteins (adjusted *p-*value < 0.05, Student’s *t*-test). Significantly down-regulated proteins are shown in green and up-regulated ones are shown in red (protein fold change| > 1.2). **(C)** Significantly changed proteins using shotgun LC-MS/MS (adjusted *p-* value < 0.05, Student’s *t*-test) from WT group and *Fmr1*^*KO*^ group were compared, and Venn diagrams demonstrate shared and unique proteins between WT group and *Fmr1*^*KO*^ group for HPT and HPC (solid line, WT group; dashed line, *Fmr1*^*KO*^ group). **(D)** Heatmaps generated using Metascape showing the significantly enriched (*p* value < 0.01) biological processes, cellular components, molecular functions and Kyoto encyclopedia of genes and genomes (KEGG) pathways for *Fmr1*^*KO*^ group in HPT and HPC. The most significant 20 terms are shown in the heatmap. Color coding indicates −log_10_ (*p* values). Rows are clustered based on their profile similarity. **(E)** Comparison of the genes from all quantified proteins and significantly changed proteins (adjusted *p-*value < 0.05, Student’s *t*-test) in HPT to gene entries from FMRP target dataset by Darnell et al (yellow), as well as from SFARI autism database (blue).

To determine proteins with significant alterations, all quantified proteins were subjected to Student’s *t*-test, and representative results for *Fmr1*^*KO*^ group in HPT and HPC were visualized in volcano plots and demonstrated in **Figure 4B** (**Supplementary Table 4**, sheet 3 and **Supplementary Table 5**, sheet 3). Points above the non-axial horizontal lines represent proteins with significant alterations (|fold change| > 1.2, adjusted *p-*value < 0.05). To explore protein alterations associated with the FXS model in HPT and HPC, we further demonstrated specific proteins which significantly changed only in WT group or *Fmr1*^*KO*^ group or showed dramatic fold changes between the two cohorts (**Figure 4C**; **Supplementary Table 6** for HPT, and **Supplementary Table 7** for HPC). To explore the macroscopic biological processes modulated by significantly altered proteins in *Fmr1*^*KO*^ group, we performed GO analysis and annotations of each cluster utilizing DAVID Bioinformatics Resources (**Supplementary Table 8**, sheet 1 for HPT WT group, sheet 2 for HPT *Fmr1*^*KO*^ group, sheet 3 for HPC WT group, and sheet 4 for HPC *Fmr1*^*KO*^ group). To better discern differential regulations in HPT and HPC, we generated heatmaps showing multi-group GO analyses in Metascape^55^. The twenty most significantly enriched biological processes, cellular components, molecular functions, and Kyoto encyclopedia of genes and genomes (KEGG) pathways are shown in **Figure 4D**, where the color-coding indicates the *p*-values of a certain term in different groups. Since more proteins were dramatically changed in HPC, most GO terms involved in HPT were also enriched in HPC, clearly showing disparities between the two regions when exposed to soy consumption. For example, we found region-specific alterations of proteins such as mitogen-activated protein kinase 1 (Mapk1), C-Jun-amino-terminal kinase-interacting protein 3, and Mbp, which demonstrated significant changes only in HPC. The subsequent gene ontology analysis also exhibits diverse biological patterns. In HPC, proteins are mostly enriched in myelin sheath, axon, and mitochondrial envelope and are associated with KEGG pathways such as synaptic vesicle cycle and glutamatergic synapse. These alterations may reflect differential responses in varied brain regions as a function of soy diet and aid the discovery of potential nutritional biomarkers. Serving as the baseline comparison, we also explored the effects of soy- and casein-based diet effects in the WT group (**Figure S6A-C**).

Furthermore, proteins that exhibited significant alterations via the Student’s *t*-test were combined and compared with the Darnell dataset as shown in the Venn diagrams for HPT (**Figure 4E**). We found 227 targets in common, of which 9 were significantly changed after soy consumption (**Supplementary Table 9**, sheet 1). We also compared our dataset with the SFARI autism database, which listed 213 genes associated with autism spectrum disorders, of which 53 were included in our quantitative data and 2 corresponded to proteins that were significantly altered in the soy cohort (**Supplementary Table 9**, sheet 1). Parallel comparisons between HPC proteins and two datasets were shown in **Figure S6B** (**Supplementary Table 9**, sheet 2).

FMRP binds to and regulates the protein synthesis of *App* mRNA, which codes for amyloid-beta precursor protein (APP)^56^. APP is processed by secretases to generate b-amyloid, the predominant peptide found in the senile plaques in AD. These findings suggest an intriguing molecular link between a neurodegenerative disorder (AD) and a neurodevelopmental syndrome (FXS).^56^ Interestingly, we found that APP was significantly decreased in HPT in the *Fmr1*^*KO*^ group (**Figure 5A**). *App* mRNA is a well-validated FMRP target that is translationally repressed by FMRP^56–57^. It remains to be determined how APP is translated and processed in response to soy consumption. To connect significantly altered proteins to genes predisposed for AD, we examined the overlap between genes linked to late-onset AD (LOAD) and proteins with altered abundance in *Fmr1*^*KO*^ brain^58^. We identified 15 significantly altered proteins genetically linked to LOAD (**Figure 5B**). 2,5-diketo-D-gluconic acid reductase B (DGKB) and progranulin (GRN) were altered in three groups, transferrin (TF) in two groups, and the remaining targets differentially expressed in one group (**Figure 5C**). These findings indicate that some of the genes linked to LOAD also have altered protein products in the *Fmr1*^*KO*^ mouse brain, further highlighting the intriguing connection between AD and FXS that warrants future mechanistic investigation.

**Figure 5.**
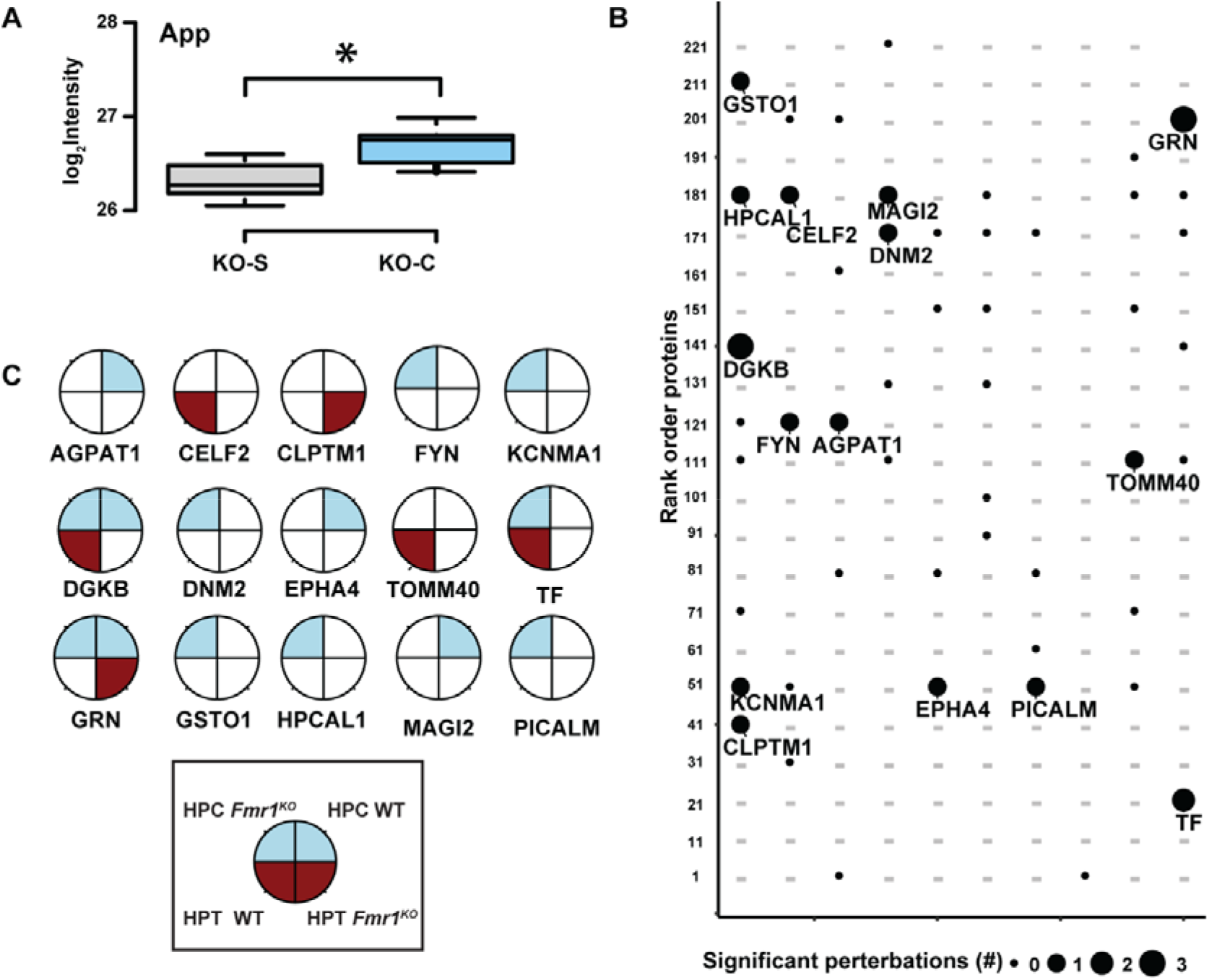
APP distribution and altered protein abundance of genes linked to LOAD. **(A)** Box plots of expression levels of APP protein between different phenotype in FXS mice in HPT. P-value < 0.05 determined by Student’s *t*-test. **(B)** Significant (FDR-adj p value < 0.05) protein changes mapped to 15 genes previously linked to LOAD (dash, not quantified). **(C)** Brain region(s) and protein abundance with significant changes plotted with brain region and genotype information. Blue, HPC; Red, HPT.

### Integrated bioinformatic analysis to decipher molecular function by combining WGCNA

This study adopted the weighted correlation network analysis (WGCNA) algorithm to better analyze and predict the molecular function of single-soy diet treatment of FXS. We were able to generate consensus networks across genotypes and phenotypes for both HPT and HPC datasets. A total of 2,108 proteins were screened for the WGCNA analysis in *Fmr1*^*KO*^ group of HPT. After normalization, no outlier samples were eliminated. The power of β◻=◻6 (scale-free R^2^◻=◻0.9) was selected as the soft-thresholding parameter to ensure a scale-free network (**Figure 6A**). A total of 19 modules with similar expression patterns were identified via average linkage hierarchical clustering (**Figure 6B**). GO analysis of the protein module members demonstrated diverse ontology for all 19 modules regarding biological functions, processes, and components (**Figure S7**). We then correlated module eigenprotein (ME), or first principal component of the module protein expression level, to different soy- or casein-based diet cohorts to assess whether a given co-expression module was related to diet (**Figure 6B**). The module and diet traits were considered statistically significant when p<0.05. We observed two modules that exhibited the strongest association with diet traits. The M5 (synapse) module, which consisted of 130 proteins, showed the strongest traits correlations with soy-based diet (corr *p* = 0.88), while the M1 (mitochondrial) module, which contained 348 proteins, exhibited the best correlation with casein-based diet (corr *p* = 0.90). By decomposing our MEs into individual diets, we assessed the relationship of MEs to different diets and measured the module eigenprotein values (summary expression profile) by case status (**Figure 6C**). We found that M1 and M5 have statistically significant difference (Z statistic and corresponding meta-analysis *p-value* < 0.05) between soy and casein cohorts. M5 (synapse) exhibited increased protein levels, while M1 (mitochondrial) showed decreased protein levels in the soy group. In addition, we performed network analysis based on the continuous measure of membership and connectivity based on WGCNA to determine the top 50 hub proteins in ME5 and ME1 and visualized using Cytoscape (**Figure 6D**). To evaluate the significance of modules, gene significance (GS) was calculated as the correlation between genes and each trait. The module membership (MM) was defined as the association between gene expression profiles and the module’s own genes. Subsequently, genes within cor.Gene MM>0.8 and cor.Gene GS>0.8 are considered key proteins. Both M1 and M5 represented the highest positive correlation with single-soy consumption (**Figure 6E**). Finally, the top 20 representative proteins were selected as key proteins in the M1 module. Rab39a, Fxyd7, and Fabp5 were upregulated in SK cohorts compared with the left one (**Figure 6F**). Differences in protein levels were assessed by Wilcoxon-signed-rank-test (\**p*-value<0.05, \*\**p*-value<0.001, \*\*\**p*-value<0.0001). WT group, as a control group, was also analyzed by WGCNA; more details showed in **Figure S8A-D** and GO analysis of the module members in **Figure S9**.

**Figure 6.**
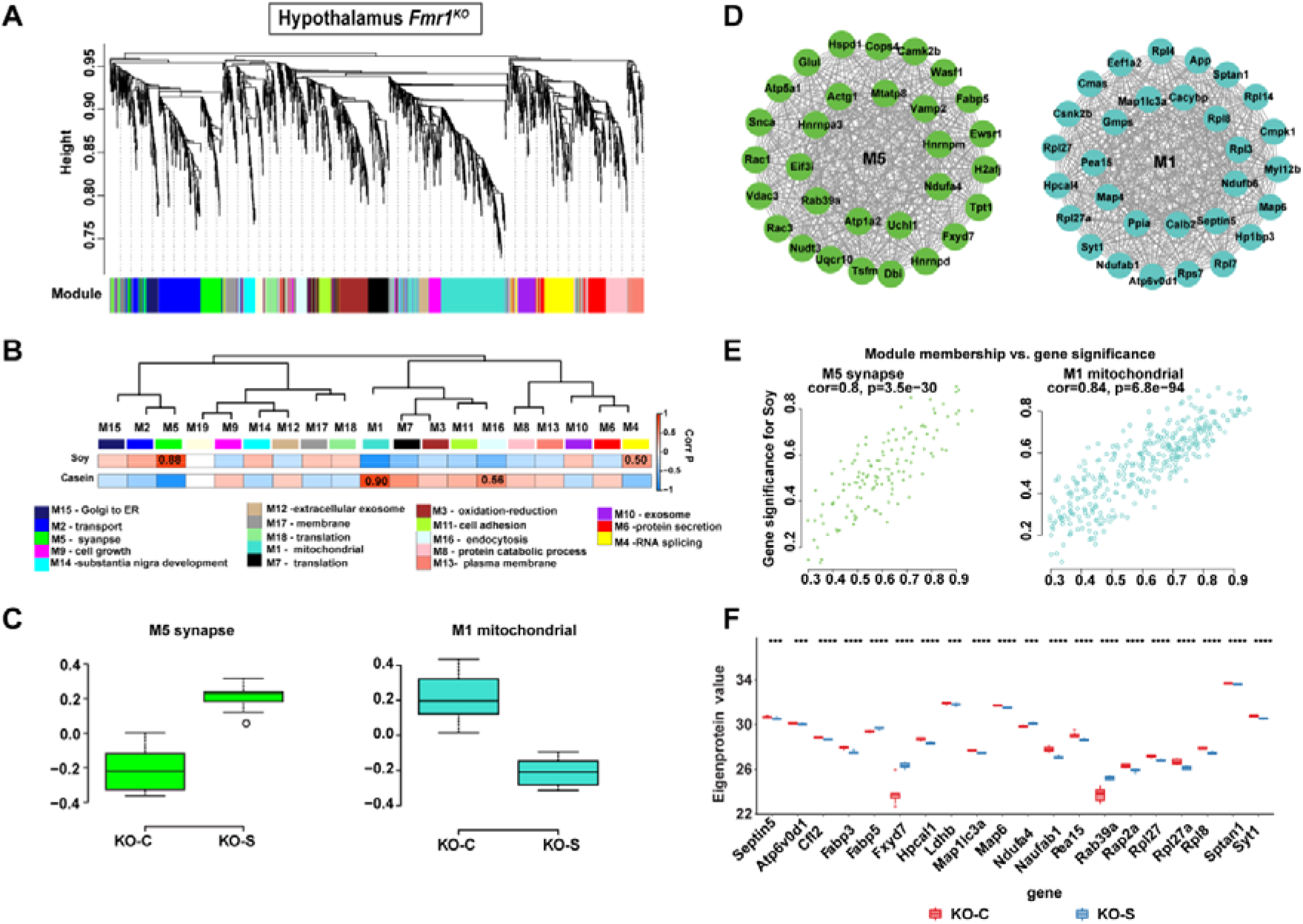
Consensus protein Co-expression Network Analysis of FXS. **(A)** Protein clustering trees generated in *Fmr1*^*KO*^ group for HPT. Each ME is in non-gray. **(B)** Protein correlation network consisting of 19 protein modules was generated from 2,108 proteins in the *Fmr1*^*KO*^ group for HPT. Module eigenproteins, which represent the first principal component of the protein expression within each module, were correlated with diet trait. Strength of positive (red) or negative (blue) correlation is shown by two-color heat map, with *P* values provided for all correlations with *P* < 0.05. Modules that showed a significant correlation with diet trait are labeled in correlation value. GO analysis of the proteins within each module clearly identified the biological functions, processes, and components associated with the module for most modules (bottom). **(C)** A synthetic eigenprotein was created to network two modules. Summary eigenprotein expression value from casein− or soy-diet for ME5 synapse (left) and ME1 mitochondrial (right) that had significant correlation to diet traits. Whiskers extend to data points that are less than 1.5* inter-quartile range away from 1^st^ and 3^rd^ quartile, respectively. The horizontal line shows the median. **(D)** Network of top 50 hub proteins in ME5 synapse (left) and ME1 mitochondrial (right) in *Fmr1*^*KO*^ group for HPT. **(E)** Visualization of gene significance (GS) vs. module membership (MM) and gene expression levels of significant modules. Scatterplot represents the GS and MM of each module that shows significant correlation (*p* < 0.05), implying that the module tends to be associated with the soy diet. Module membership versus gene significance for ME5 synapse (left) and ME1 mitochondrial (right). **(F)** Top 20 representative proteins selected in ME1 mitochondrial. Box plots represent the median, 25th and 75th percentiles, and whiskers represent measurements to the 5th and 95th percentiles. Differences in protein levels were assessed by Wilcoxon-test (\**p*-value<0.05, \*\**p*-value<0.001, \*\*\**p*-value<0.0001).

To assess whether the network was similar in other brain regions, we conducted the co-expression network analysis for HPC *Fmr1*^*KO*^ group (**Figure S10A-D**). A total of 2,061 proteins were used to generate 6 protein co-expression modules, which ranged in size from 662 proteins (M1) to 69 proteins (M6). GO analysis of the proteins within each module clearly identified the biological processes associated with the module. Other GO analyses of cellular components and molecular function terms are listed in **Figure S11**. We observed three modules that were significantly correlated with diet phenotype (see **Figure S10A**, right panel): modules M1 mitochondrial, M2 synapse, and M3 metabolism. The M3 metabolism module showed the strongest single-soy diet trait correlation (corr *p* = 0.90), but M1 and M2 module showed the strongest negative correlation of soy trait (M1 corr *p* = 0.90; M2 corr *p* = 0.90). Noticeably, M1 and M2 modules were decreased in the KO-S cohort compared to the KO-C cohort, but the M3 module showed the opposite trend (**Figure S10B**). In addition, we performed network analysis based on the continuous measure of membership and connectivity based on WGCNA to determine the top 50 hub proteins in ME1, ME2, and ME3 and visualized using Cytoscape (**Figure S10C**). Finally, the top 20 representative proteins (cor.Gene MM>0.8 and cor.Gene GS>0.8) were selected as key proteins in M3 module. Differences in protein levels were assessed by Wilcoxon-signed-rank-test (\**p*-value<0.05, \*\**p*-value<0.001, \*\*\**p*-value<0.0001, see **Figure S10D**). Most proteins were decreased in KO-S cohort compared to KO-C cohort, except Srsf3, Phb, Nrgn, Mp68, Gng7, and Cox6a1. WT group as a control group was also analyzed by WGCNA with details shown in **Figure S12A-C** and GO analysis of the module members in **Figure S13**.

Noticeably, the module function of mitochondrion-related module eigenproteins values by case status showed similar differences between case cohorts in HPT and HPC. Recent work by Licznerski et al. suggests that loss of FMRP causes a mitochondrial inner membrane proton leak that prevent synaptic maturation^59^. In order to explore the effect of soy consumption and *Fmr1* genotype on mitochondrial regulation, we compared the proteins included in the mitochondrial module with glycolytic enzymes and with enzymes required for the TCA cycle (**Supplementary Table 10**). We found that those enzymes involved in glycolytic and TCA cycle pathways were dysregulated by soy consumption in both HPT and HPC (**Figure S14A**). These findings suggest a consensus in altered metabolic enzyme levels across brain regions in response to soy consumption. Nonetheless, the synapse module exhibited opposite effects in HPT and HPC, indicating complex proteome expression profiles across brain regions.

### Spatially resolved biomarkers correlated with shotgun proteomics analysis

We also compared the peptide list from MALDI MSI experiments to all quantified peptides and significantly changed peptides revealed by LC-MS/MS in WT group and *Fmr1*^*KO*^ group in the HPT (**Figure 7A**) and the HPC (see **Figure S14B**). The LC-MS/MS proteomics data had an overlap of an average of 6% of the imaging proteins. Among the common proteins, Atp1a2 exhibited the same dysregulated trend by both LC-MS/MS and spatially resolved proteomics. The Atp1a2 gene is part of the seizure susceptibility locus in mouse^60^. Regardless of the *Fmr1* genotype, Atp1a2 protein expression increased with soy diet, but the finding was more pronounced in *Fmr1*^*KO*^ than WT HPT (**Figure 7B**). The ROC plot also showed that the AUC value was higher in *Fmr1*^*KO*^ (**Figure S14 C**). In addition, we found that stathmin (Stmn1), which is involved in MAPK signaling, exhibited significant upregulation in the HPC in *Fmr1*^*KO*^ mice in response to soy diet (**Figure 7C**), suggesting a potential role in cognition. Other well-known FMRP targets resolved in spatial proteomics and shotgun proteomics that were significantly changed include Syn1, Aldoa, Mbp, and other new candidates (**Supplementary Table 11**). Noticeably, Gapdh, a key glycolytic enzyme (listed in **Supplementary Table 9**) that is associated with an elevated risk of developing AD^61^,was only identified in the imaging data pool. Gapdh was upregulated in the soy cohorts compared to the casein cohorts across brain regions in *Fmr1*^*KO*^ group (**Figure 7D**). The same result was found in WT group (**Figure S14D**). Thus, consumption of a single source soy protein-based diet is associated with increased expression of Gapdh in both HPT and HPC in WT and *Fmr1*^*KO*^ mice, albeit the effect was more pronounced in the *Fmr1*^*KO*^. The mechanism and biological significance of elevated Gapdh levels for FXS remains to be determined. In addition, metabotropic glutamate receptor 5 (mGluR5) was significantly down-regulated in *Fmr1*^*KO*^ group in HPC in response to soy diet (see **Figure 7E**, p-adjust value<0.05, log_2_(fold change) = −1.5) but notably not in WT group. Multiple studies have shown that altered mGluR5 signaling plays an important role in FXS pathophysiology^62^. These findings further highlight the potential impact of the soy diet on neural plasticity and behavioral responses. Taken together, all quantified proteins are highly correlated with synapse and mitochondrial energy module functions.

**Figure 7.**
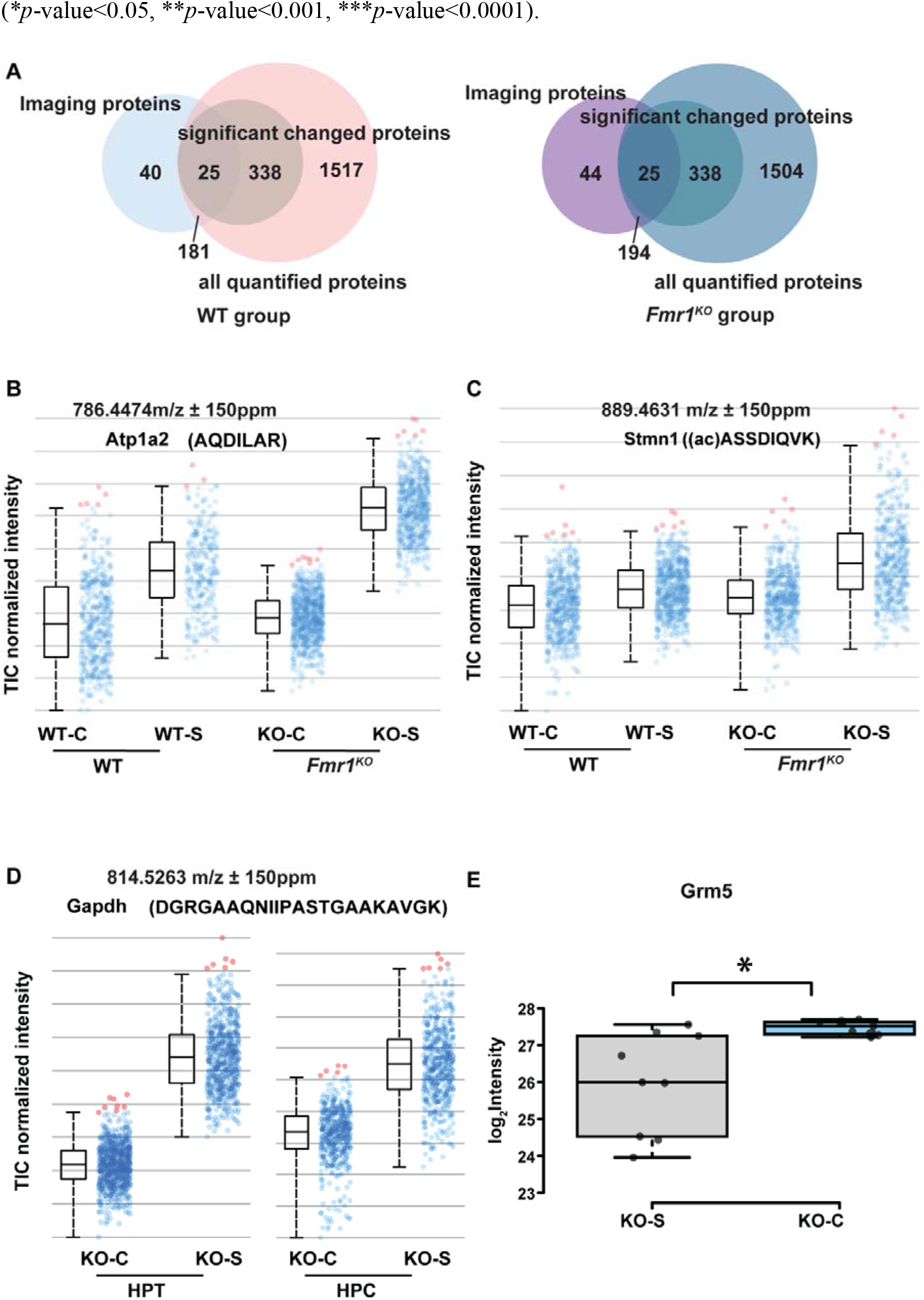
Spatially resolved candidate protein biomarkers of *Fmr1*^*KO*^. **(A)** Venn diagram showing the overlap among annotated imaging proteins, LC-MS/MS all quantified protein numbers, and significantly changed proteins in WT group (left panel) and *Fmr1*^*KO*^ group (right panel) in HPT. **(B)** Normalized intensity of total ion signals of the Atp1a2 protonated peptide in four cohorts. The boxplot points correspond to relative intensities of *m/z* 789.4474-related peaks taken from each spectrum in all images acquired. **(C)** Normalized intensity of total ion signals of the Stmn1 protonated peptide in four cohorts. The boxplot points correspond to relative intensities of *m/z* 889.4631-related peaks taken from each spectrum in all images acquired. **(D)** Normalized intensity of total ion signals of the Gapdh protonated peptide in four cohorts in HPT (left panel) and HPC (right panel) of *Fmr1*^*KO*^ group. The boxplot points correspond to relative intensities of *m/z* 814.5263-related peaks taken from each spectrum in all images acquired. **(E)** Box plots of expression levels of Grm5 protein between different phenotype in FXS mice in HPC. P-value < 0.05 determined by Student’s *t*-test.

## Discussion

Consumption of single-source, soy-based diets are associated with increased seizures, body weight, and autistic behavior in neurodevelopmental disability models^7, 10,11–12^. Even though proteomics studies in FXS mouse model have been performed in cell culture (SILAC-Labeled, iTRAQ-labeled), and blood (de novo protein)^63–64^, few studies investigate potential correlation of molecular biomarkers with specific diets, including soy-based diets, which impedes the understanding of how diet affects neurological development and outcomes. In this study, state-of-the-art spatially resolved proteomic MALDI-MS imaging was employed and LC-ESI MS/MS quantitative proteomics was integrated to systematically evaluate the spatial and dynamic changes of candidate protein and peptide biomarkers induced by consumption of soy protein-based diet by *Fmr1*^*KO*^ mouse. We successfully develop a high sensitivity and robust strategy to resolve the spatial distribution of protein candidates, which identified 1640 mass spectral peaks across whole brain region of four different cohorts (**Figure S1B**). To better understand the relationship between aberrant protein spatial distribution and FXS pathology of soy consumption, MS imaging peaks were further annotated based on on-tissue peptide extraction results. Nearly 90% of unique peptides from LC-MS/MS could be matched with the MS imaging peak list, and at least 66% of the unique sequences in each cohort could be annotated in two or more biological replicates where only [H]^+^ adducts were considered within mass error tolerances less than 150 ppm. Of note, 1,004 protonated peptides, corresponding to 350 proteins were assigned based on MS^2^ unique sequences. Thus, the spatially resolved proteomics strategy, guided by MALDI-MSI tryptic digested peptide imaging, could potentially allow mapping of the global-scale peptide changes in FXS. Interestingly, peptide identification in the KO-S cohort involved targets implicated in autism, transport function, and synaptic transmission. In particular, the altered regulation of Eef2 in both HPT and HPC in *Fmr1*^*KO*^ mice, suggests that mGluR5 signaling plays an important role in the *Fmr1*^*KO*^ response to soy consumption.

To overcome the challenges posed by lower S/N of MALDI-produced precursor ions and limited quantification capacity, we conducted LC-MS/MS label-free quantitative analysis of proteomes extracted from isolated HPC and HPT brain regions. Overall, we quantified 82 (WT group), 62 (*Fmr1*^*KO*^ group) proteins in HPT and 428 (WT group), 730 (*Fmr1*^*KO*^ group) proteins in HPC that were significantly and differentially expressed as a function of *Fmr1* genotype and soy protein-based diet. Many played critical roles in the postsynaptic structure and signaling among our list of significantly altered proteins in *Fmr1*^*KO*^ group in HPT and HPC. Interestingly, most GO terms involved in HPT were also enriched in HPC, clearly showing disparities between the two regions when exposed to soy consumption. For example, Mapk1 and Mbp demonstrated significant changes only in HPC, while APP exhibited significant changes in HPT of the *Fmr1*^*KO*^ group. These alterations may indicate that varied brain regions differentially respond to soy diet. We further connected proteins to genes that predispose for AD. Our screen identified 15 significantly altered proteins genetically linked to LOAD, such as progranulin (*GRN*), which mediates neuroinflammatory function related to multiple neurodegenerative diseases^65^, such as Parkinson’s disease, AD, and amyotrophic lateral sclerosis, and is also dysregulated in HPT *Fmr1*^*KO*^ group. These findings indicate that some of the genes linked to LOAD also have altered protein products in the *Fmr1*^*KO*^ mouse brain, further highlighting the intriguing connection between AD and FXS, which warrants future mechanistic investigation. To arrive at a consensus view of the proteomic changes in brain HPT and HPC region response to soy consumption in WT and *Fmr1*^*KO*^ mice, we performed WGCNA analyses and revealed that the protein co-expression families most strongly correlated to the soy diet included synaptic and mitochondrial families. Decreased key glycolytic and tricarboxylic acid (TCA) cycle enzyme levels, which could explain abnormal mitochondrial proton leak that impair synaptic plasticity^59^, may be regulated by soy consumption. Finally, when comparing shotgun proteomic results with MALDI-MSI data, we found that LC-MS/MS proteomics data were mapped to approximately 6% of proteins found in MS imaging datasets. Of note, some well-known FMRP targets, like Eef2, Atp1a2, and Stmn1, showed overlapping spatial distribution and in LC-MS/MS quantification results. However, there still were some unique protein candidates only detected in one strategy. Gapdh, a key glycolytic enzyme, was upregulated in the soy cohorts as compared to the casein cohorts across the brain region in *Fmr1*^*KO*^ group, in the imaging protein candidate pool. mGluR5, which is commonly linked to FXS pathology, was down-regulated in *Fmr1*^*KO*^ group in HPC in the LC-MS/MS data.

Overall, the novel, state-of-the-art spatially resolved proteomics methodology employed here has strong potential to identify protein biomarkers in response to genetics, diet, and drug treatment. This procedure may be applied to various diseases to identify diagnostic or prognostic indicators and to further identify targets for therapeutic intervention.

## Methods

### Mouse husbandry, test diets, and tissue collection

Wild type (WT) and *Fmr1*^*KO*^(KO) mice in the C57BL/6J background were bred in our colony maintained at the University of Wisconsin-Madison. Animals were housed in microisolator cages on a 12-hour light cycle with *ad libitum* access to food (Teklad 2019) and water. All husbandry and euthanasia procedures were performed in accordance with NIH and an approved University of Wisconsin-Madison Institutional Animal Care and Use Committee (IACUC) protocol administered through the University of Wisconsin Research Animal Resource Center. Breeder *Fmr1*^*HET*^ females and WT and *Fmr1*^*KO*^ males were transferred to the test diets for at least 12 days prior to breeding. Test diets were formulated and synthesized by Envigo Teklad Diets (Madison, WI) based on their AIN-93G diet (TD.94045). The soy protein-based diet (TD.180375) contained soy protein isolate swapped for casein protein and matched for macronutrient content (protein 18.8% kcal, carbohydrate 64.1% kcal, fat 17.1% kcal; 3.8 kcal/g) and included 0.5% calcium, 0.3% available phosphorus, 0.2% sodium, 0.36% potassium, 0.3% chloride and 0.05% magnesium. Green food dye was added for visual differentiation. The casein protein-based diet (TD.180374) was a modified version of AIN-93G (TD.94045) with the sodium increased to 0.2% to match the soy diet and red food dye added for visual differentiation (protein 18.9% kcal, carbohydrate 63.8% kcal, fat 17.3% kcal; 3.7 kcal/g). Offspring (WT and *Fmr1*^*KO*^ male littermates) were weaned at postnatal day 18 (P18) and maintained on their respective diets until tissue harvest at 3 months. Genotypes were determined by PCR analysis of DNA extracted from tail biopsies taken at weaning. For tissue harvest, mice were anesthetized with isoflurane, perfused with 30 mL phosphate-buffered saline via the left ventricle, decapitated, and either: (1) brain regions dissected including HPT and HPC and tissue flash frozen on dry ice and stored at −80°C, or (2) the whole intact brain removed from the skull, cut in half along the midline with hemispheres immediately placed in disposable base molds and covered/embedded with melted 10% gelatin. Gelatin molds were stored at −80°C until processing. Eight-μm thick sagittal tissue sections were obtained using a cryostat microtome (HM525, Thermo Fisher Scientific, Waltham, MA). Three consecutive sections from each brain were thaw-mounted onto ITO-coated glass slides and stored at −80 °C. All sections underwent the tissue digestion procedure, with one section used for MADLI-IMS and two sections used for annotation experiments by LC-MS/MS. For global quantitative proteomic analyses, HPT and HPC were sonicated (3-s on and 3-s off, amplitude 50%) and subjected to enzymatic digestion.

### Sample preparation for MALDI MS imaging

Endogenous background compounds (lipids, peptides) were removed using original washing or Carnoy’s washing methods^66^. In original method, tissues were washed twice in 70% ethanol for 30 sec and 100% ethanol for 30 sec. In Carnoy’s method, tissues were washed as follows: submerge in (i) 70% ethanol for 30 sec; (ii) 100% ethanol for 30 sec; (iii) Carnoy’s buffer (ethanol: chloroform: acetic acid 6:3:1) for 2 min; (iv) 100% ethanol for 30 sec; (v) Optima grade water for 30 sec; and (vi) 100% ethanol for 30 sec. The sections were then allowed to dry for 15 min under vacuum. Enzyme and matrix application were performed by a robotic TM sprayer system (HTX Technologies, Carrobo, NC). For enzyme deposition, trypsin was dissolved in 15 mM ammonium bicarbonate to a final concentration of 0.05 μg/μL, sprayed at a flow rate of 0.02 mL/min a total of 8 passes performed. The nozzle temperature was set to 30°C, with a moving velocity of 750 mm/min. Then, tissue sections were incubated in a humidity chamber at 37°C for 4 hr. CHCA dissolved in 70% acetonitrile (ACN), 1% TFA solution at a concentration of 10 mg/mL was used as the matrix for peptide imaging. A total of 4 passes of matrix spraying were performed at a flow rate of 0.12 mL/min, and 30 s drying time between each pass. The nozzle temperature was set to 75°C with a moving velocity of 800 mm/min. The slides were dried in a desiccator at room temperature for 15 min and subjected to instrument analysis.

### MALDI MS imaging

MALDI-MS imaging was performed on a Bruker RapifleX MALDI Tissuetyper TOF/TOF (Bruker, Billerica, MA) equipped with a smart-beam 3D 10 kHz laser. Samples were analyzed in positive-ion mode with an m/z range of 600 to 3000, 80% laser energy, and 50 μm step size.

### Peptide extraction

Consecutive tissue sections were prepared for MALDI-MSI using the same sample preparation protocol as above. After trypsin deposition, the proteolytic peptides were extracted from the ITO slides using following protocol: (i) 40 μL of 0.1% TFA (repeat 4 times); (ii) 40 μL of 50% ACN, 0.1% TFA (repeat 4 times); (iii) 90% ACN, 0.1% TFA (repeat 4 times). Four extracts from each brain were combined, dried, and resuspended in 0.1% TFA and then cleaned with Omix C18 tips (Agilent). The purified extracts were dried and stored at −20 °C until LC-ESI-MS/MS analysis.

### Sample preparation for global proteomics analysis

HPT and HPC tissues were homogenized in lysis buffer (8◻M Urea, 50◻mM tris) using a probe sonicator, and protein concentration was determined using a BCA Protein Assay Kit (Thermo Pierce, Rockford, IL). Samples were reduced with 100 mM DTT for 1hr, alkylated with 200 mM IAA for 30 min before quenching with 100 mM DTT. Proteins were digested by trypsin at 37°C for 16 hr in a 50:1 (protein:enzyme) ratio. Digests were quenched by lowering the pH to < 3 with 10% TFA. Peptides were desalted with SepPak C18 solid-phase extraction (SPE) cartridges (Waters, Milford, MA). All samples were dried in vacuo and stored at −80°C until LC-ESI-MS/MS analysis.

### LC-MS/MS acquisition

Samples from imaging peptide extraction were dissolved in 0.1% FA and analyzed on the Orbitrap Fusion™ Lumos™ Tribrid™ Mass Spectrometer (Thermo Fisher Scientific, Bremen, Germany) coupled to a Dionex UPLC system. Samples global proteomics analysis were analyzed on the Orbitrap Q-Exactive HF Mass Spectrometer (Thermo Fisher Scientific, Bremen, Germany). The chromatographic separation was carried out via mobile phase A that consists of 0.1% FA in water and mobile phase B consisting of 0.1% FA in ACN. Peptides were loaded onto a 75 μm × 15 cm homemade column packed with 1.7 μm, 130 Å, BEH C18 material obtained from a Waters UPLC column (part no. 186004661). Emitter tips were pulled from capillary tubing 75 um id (Polymicro Technologies, Phoenix, AZ) using a model P-2000 laser puller (Sutter Instrument Co., Novato, CA).

For imaging peptide extracts, the LC gradient was set as follows, 5%-8% B (18-30 min), 8%−22% B (30-100 min) and 22%−40% B (100-120 min) with a flow rate of 300 nL/min. Survey scans of peptide precursors were performed with a scan range from 300 to 1500 *m/z* at a resolving power of 60 K (at *m/z* 200) with automatic gain control (AGC) target of 2 × 10^6^ and a maximum injection time of 100 msec. The top 20 precursors were then selected for higher-energy C-trap dissociation tandem mass spectrometry (HCD MS^2^) analysis with an isolation width of 1 Da, a normalized collision energy (NCE) of 30, a resolving power of 15 K (at *m/z* 200), an AGC target of 1 × 10^4^, a maximum injection time of 100 msec, and a lower mass limit of 120 *m/z*. Precursors were subject to dynamic exclusion for 45 sec with a 10 ppm tolerance. Each sample was acquired in technical triplicate.

For global proteomics samples, the LC gradient was set as follows: 3%-10% B (18-31 min), 10%-24% B (31-95 min) and 24%-35% B (95-127 min) with a flow rate of 300 nL/min. Survey scans of peptide precursors were performed with a scan range from 300 to 2000 *m/z* at a resolving power of 60 K (at *m/z* 200) with an AGC target of 1 × 10^6^ and a maximum injection time of 100 msec. The top 15 precursors were then selected for HCD MS^2^ analysis with an isolation width of 2 Da, a normalized collision energy (NCE) of 28, a resolving power of 15 K (at *m/z* 200), an AGC target of 2 × 10^5^, a maximum injection time of 150 msec, and a lower mass limit of 100 *m/z*. Precursors were subject to dynamic exclusion for 18 sec with a 10-ppm tolerance. Each sample was acquired in technical triplicate.

### Data analysis

For MALDI MS imaging data analysis, regions of interest ROIs were selected in flexImaging (Bruker, Billerica, MA) and imported into SCiLS Lab for spectra smoothed alignment^42^, and total ion intensity (TIC) normalization. Overview spectra were then exported for peak picking to mMass and peptide annotation was performed by matching tissue-specific peaks with on tissue peptide extraction results from LC-MS/MS. The detected peaks were re-imported into SCiLS Lab for subsequent segmentation and statistical analysis. t-SNE clustering was accomplished using a python implementation of the t-distributed Stochastic Neighbor Embedding algorithm, which is made available through the machine learning library, scikit learn. Iterative optimization was performed to establish the most appropriate parameters.

Protein identification and quantification by MaxQuant (version 1.5.3.8) based database searching, using the integrated Andromeda search engine with FDR < 1% at peptide and protein levels. The tandem mass spectra were searched against the *Mus musculus* UniProt database (version updated December 2018). A reverse database for the decoy search was generated automatically in MaxQuant. Enzyme specificity was set to ‘Trypsin/p’, and a minimum number of seven amino acids were required for peptide identification. For label-free protein quantification (LFQ), the MaxQuant LFQ algorithm was used to quantitate the MS signals, and the proteins’ intensities were represented in LFQintesnity^45^. Cysteine carbamidomethylation was set as the fixed modification. The oxidation of M and acetylation of the protein N-terminal were set as the variable modifications. The first search mass tolerance was 20 ppm, and the main search peptide tolerance was 4.5 ppm. The false discovery rates of the peptide-spectrum matches, and proteins were set to less than 1%. For peptide quantification, the intensities of all samples were extracted from the MaxQuant result peptide files. Then, the expression matrix was subjected to normalization followed by log2-transformed by Perseus^54^. Bioinformatic analyses was performed with R software environment. GO analyses were generated using Metascape^55^ (version 3.5) and DAVID bioinformatics resources^67^ with a FDR cutoff of 0.05. WGCNA algorithm was used for network analysis as previously described^68^.

## Data availability

The mass spectrometry data have been deposited to the ProteomeXchange Consortium^69^ via the PRIDE^70^ partner repository with the dataset identifier PXD028146. Public release of the data will be made on time for online publication of the paper.

## Code availability

All data and files needed to reproduce t-SNE clustering have been included in the supplemental information and may also be found online https://github.com/grahamdelafield/soy_casein_mouse_tSNE.

## Acknowledgements

This work was supported by the United States Department of Agriculture (grant number 2018-67001-28266 to CW and LL) and the University of Wisconsin Pandemic Affected Research Continuation Initiative (PARCI). L.L. would like to acknowledge NIH grants R01DK071801, R56MH110215, RF1AG052324, NCRR S10RR029531, and S10OD025084, as well as a Vilas Distinguished Achievement Professorship and Charles Melbourne Johnson Professorship with funding provided by the Wisconsin Alumni Research Foundation and University of Wisconsin-Madison School of Pharmacy. MX would like to acknowledge the National Science Foundation Graduate Research Fellowship Program under Grant No. DGE-1747503. The funding bodies played no role in study design; the collection, analysis, and interpretation of data; manuscript preparation; or the decision to publish.

## Author contributions

M.M. and Q.Y. contributed equally to this work. M.M., Q.Y., C.W., and L.L. designed the study and conceived the experiments. M.M. and Q.Y. conducted the experiments, analyzed data, and wrote the first draft of manuscript; G.D., Y.C., Z.L., W.X., M.X. assisted with data analysis and organization and some figure making; X.S. did the H&E staining; A.J and P.W. breed the mice and gelatin embedded. M.M., Q.Y., C.W., and L.L. prepared and revised the manuscript and all authors provided editorial feedback and input.

