## supplementary figure for "On-tissue spatial proteomics integrating MALDI-MS imaging with shotgun proteomics reveals soy consumption-induced biomarkers in a fragile X syndrome mouse model"

**t-SNE Analysis.**

Prior to clustering, mouse brain regions were annotated from stained brain sections, which were used as a reference to define hippocampus and hypothalamus regions within the MALDI data sets. Using SCiLS software, all brain sections were loaded into a single SCiLS analysis, and the tissue-level total ion count was used as the intersection normalization factor. Once signal intensities were normalized, polygons were drawn around the hippocampus and hypothalamus brain regions. All pixels within these drawn polygons were exported for clustering.

Leveraging the Python library PyimzML, hippocampus and hypothalamus pixels were concatenated into individual two-dimensional arrays with each pixel assigned an integer identification code that corresponded to treatment group (Supplemental Figure 4 A). These two-dimensional arrays serve to correlate pixels to treatment group identity after clustering was performed. We then extracted the pixel-level intensity of each m/z value that was assigned to a tryptic peptide identity, resulting in a *N x n x M* data array (Supplemental Figure 4 B. N = number of tissues, n = number of pixels in each tissue, M = number of masses extracted). This data array was then collapsed into two dimensions by linearizing all data belonging to a single m/z value (Supplemental Figure 4 C). This final array was used as the test set for t-SNE clustering.

t-SNE clustering was accomplished using a python implementation of the t-distributed Stochastic Neighbor Embedding algorithm, which is made available through the machine learning library, scikit learn. Iterative optimization was performed to establish the most appropriate parameters. Of note, perplexity and number of clustering iterations were the parameters of greatest effect, ultimately showing that a perplexity=50.0 and iterations=4,000 yielded the best results. Resulting data were plotted in three dimensions, which may be viewed interactively online (<https://github.com/grahamdelafield/soy_casein_mouse_tSNE/tree/main/Figures>).

**Supplementary Figures**

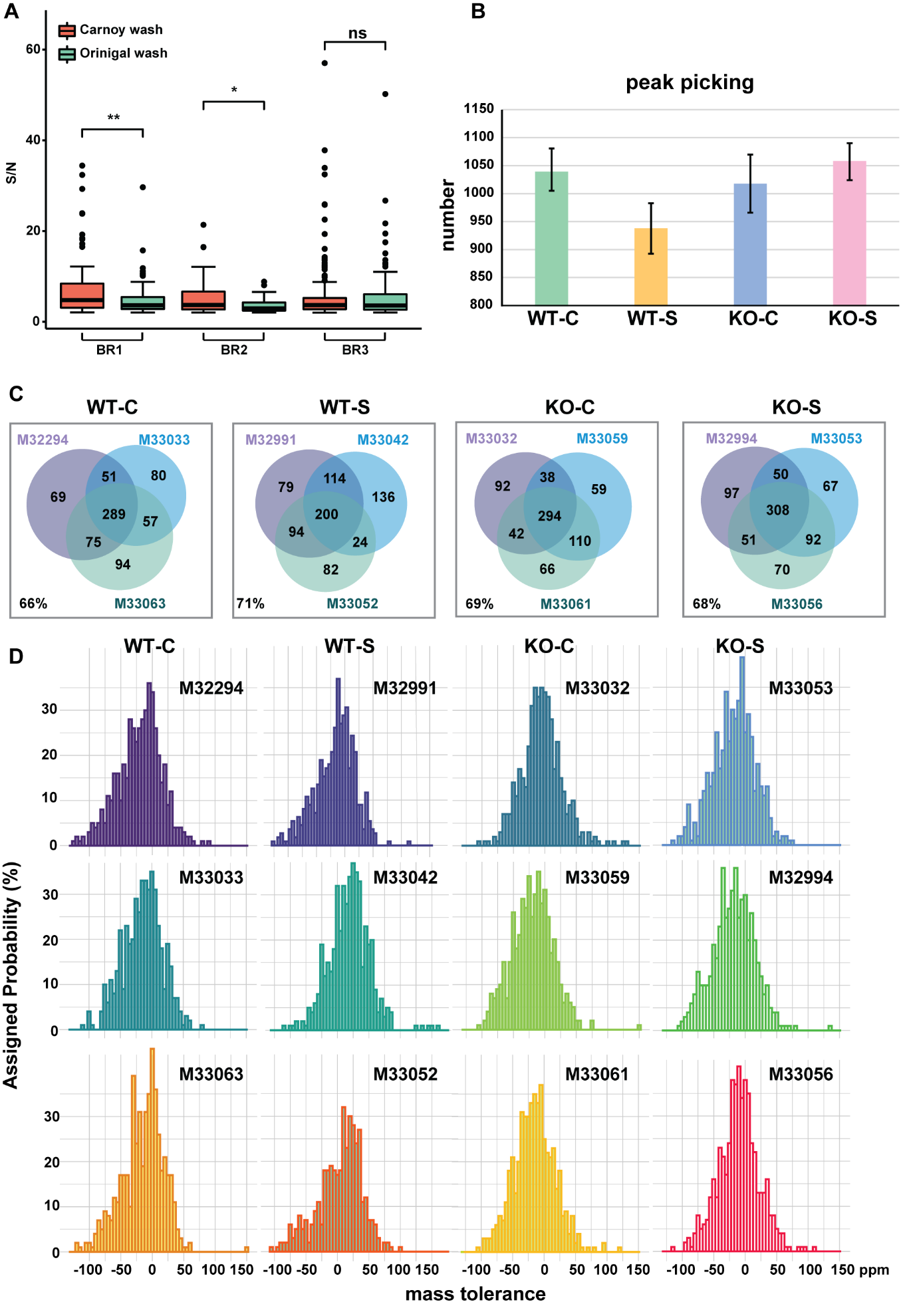

**Figure S1. MALDI-MSI data reproducibility.**

(A) Distribution of S/N of peaks detected in different mass ranges across the whole brain region, comparing Carnoy washing method with original washing method of three biological replicates (BR) (BR1-BR3). The significance of S/N between different washing method was assessed by Wilcoxon signed-rank test (**p*-value<0.05, ***p*-value<0.001, ns non-significance).

(B) Bar graph shows statistical test of the number of peak pickings based on Figure 1B strategy of four cohorts. Each cohort has three biological replicates, N=3.

(C) Venn diagram of annotated, protonated, unique peptides overlap among three biological replicates of each cohort, with each identified by assigned mouse ID number. The percent of peptides, which could be identified in at least two biological replicates, are labeled in the bottom left inside each rectangle.

(D) Histograms showing the average frequency mass tolerance of on-tissue imaging, peptide annotation of each mouse.

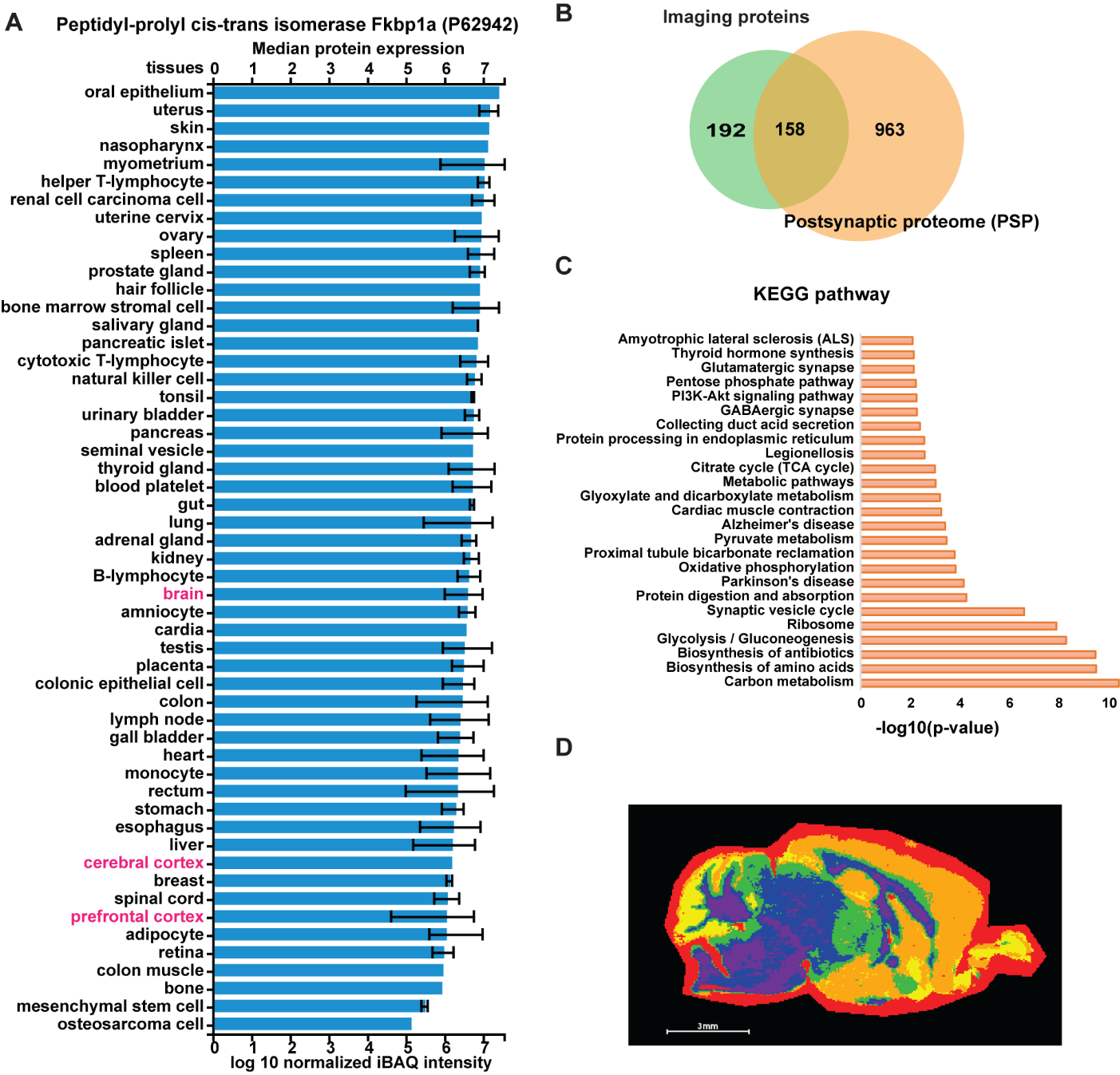

**Figure S2. MSI gene ontology and segmentation analysis.**

(A) Distribution of Fkbp1a protein in human tissues from MaxQB database^1^. Human brain related tissues are highlight with red color.

(B) Comparison of the proteins identified from MS imaging workflow (light green) with mouse brain synaptic proteins (peach) (G2Cdb:PSP).

(C) KEGG pathway enrichment analysis of 350 proteins by Fisher’s exact test. Top 25 most significant pathways were plotted with the x axis representing significance values.

(D) Histological annotation and segmentation analysis using the Bisecting k-Means and correlation distance approach.

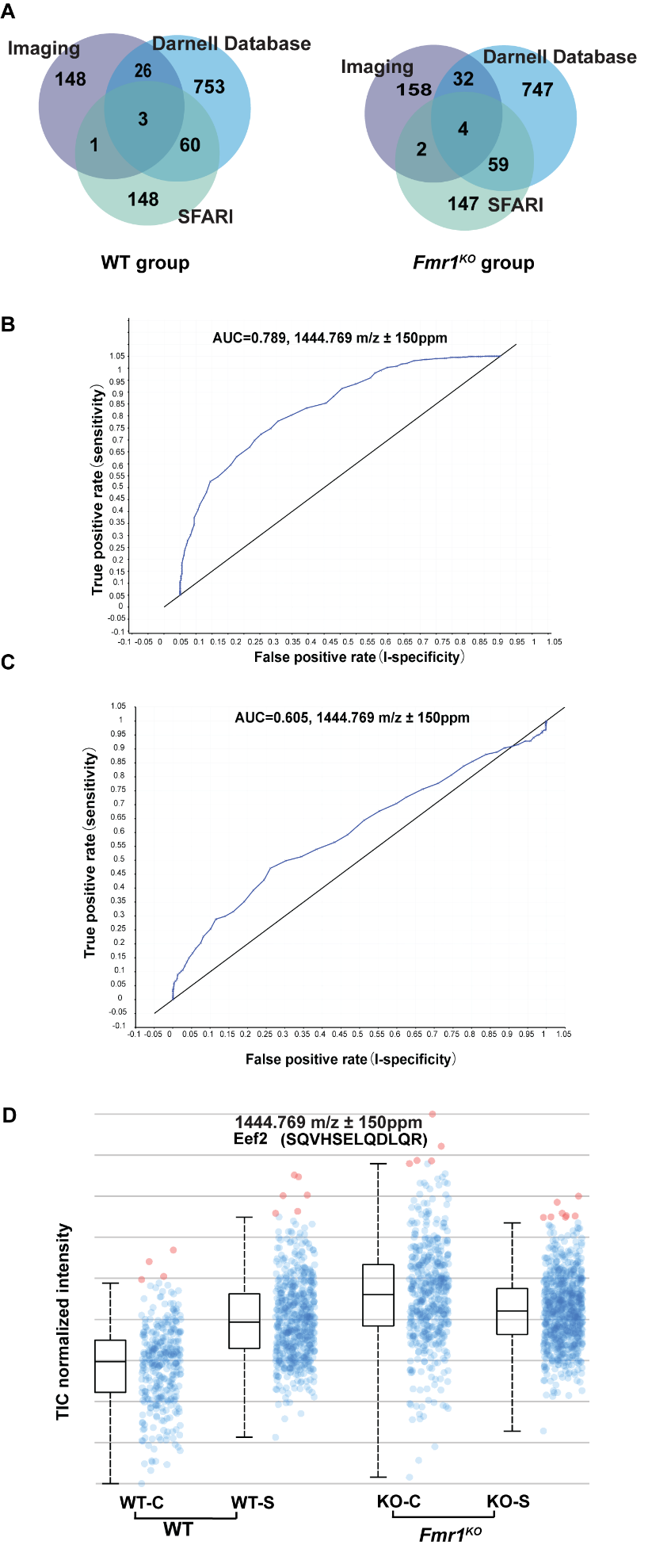

**Figure S3. MSI data correlation with public database.**

(A) Comparison of the proteins identified from MS imaging workflow (purple) with the SFARI autism database (light green), as well as from FMRP target database by Darnell et al^2^ (blue).

(B) Adjusted receiver operating characteristic (ROC) analysis corresponding to Figure 3B in WT group of HPC.

(C) Adjusted ROC analysis corresponding to Figure 3B in *Fmr1^KO^* of HPC.

(D) Normalized intensity of total ion signals of the Eef2 protonated peptide in four cohorts in HPT. The boxplot points correspond to relative intensities of *m/z* 144.769-related peaks taken from each spectrum in all images acquired.

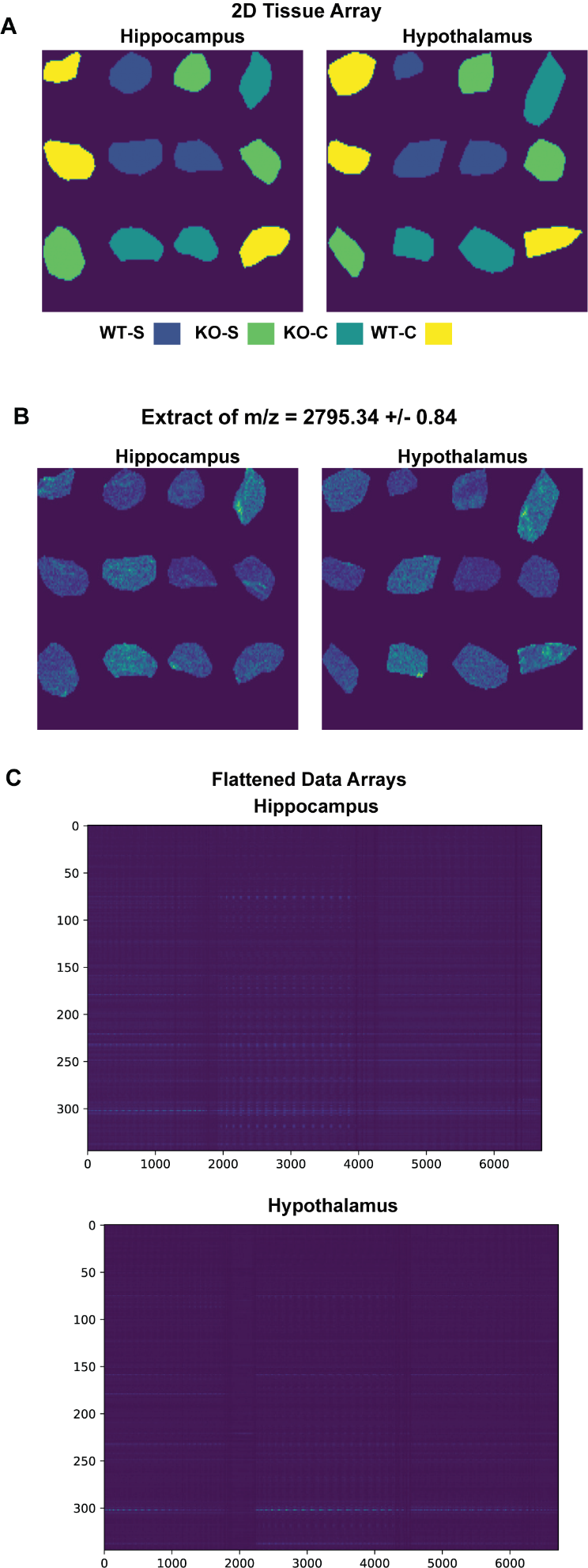

**Figure S4. t-SNE analysis.**

(A) Two dimensional arrays representing the outlined brain regions exported from SCiLS software. Each pixel is color coded according to treatment group and every pixel within individual polygons has been supplied a unique integer identification. These identification codes are used to correlate pixels to treatment group identity after clustering is performed.

(B) Data arrays containing the pixel-level intensity of *m/z* = 2759.34. These ion images were constructed for each *m/z* that was assigned to a tryptic peptide identification and compiled into a large three-dimensional master array.

(C) Flattened data arrays containing all pixel information from all hippocampus and hypothalamus brain regions. These arrays are constructed by linearizing the pixel information from the 3D master array. Viewing these heatmaps, each row represents a unique *m/z* value (i.e. tryptic peptide identification) and each column represents a unique pixel.

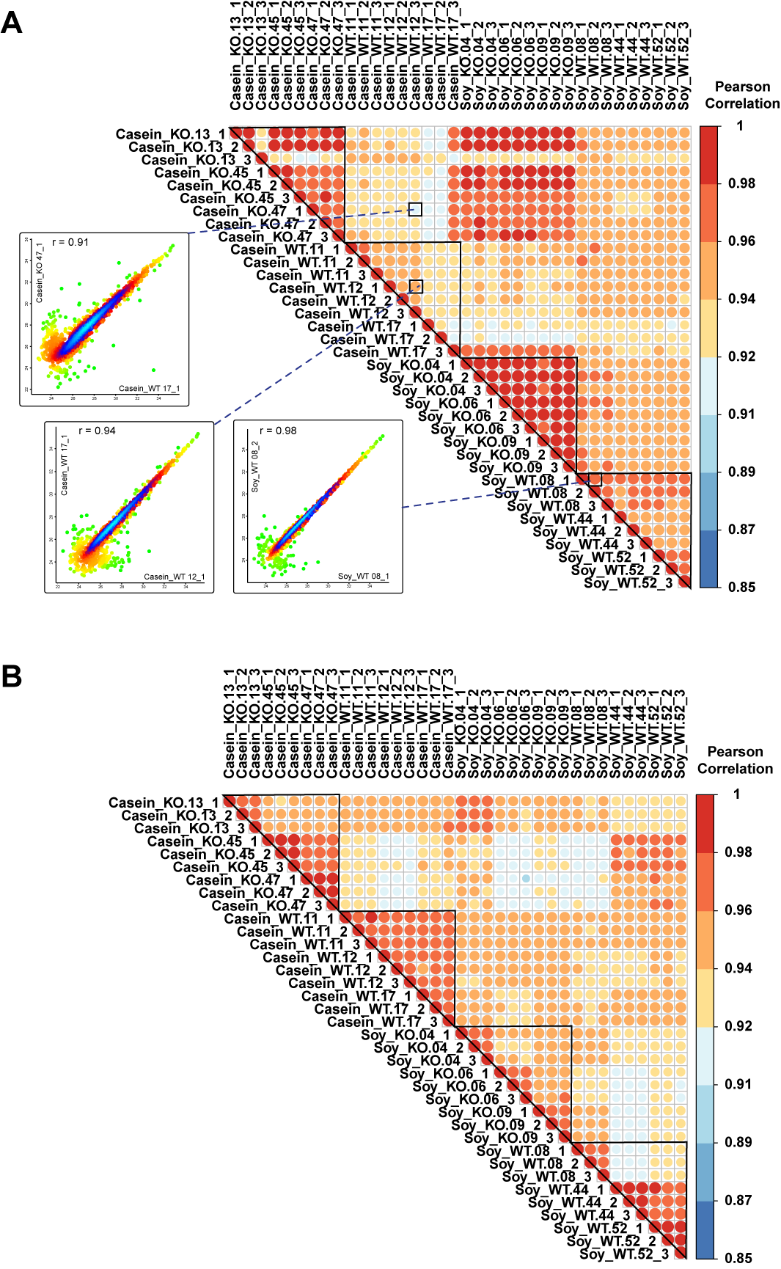

**Figure S5. Shotgun Proteomics LC-MS/MS Data Reproducibility.**

(A) Pearson correlation heatmap of quantified protein intensities in all technical replicates (n=3) and biological replicates (n=3) from 36 LC-MS/MS runs for HPT. Density plots illustrate the protein intensity correlation of two representative pairs comparing technical replicates (lower right inset), biological replicates (lower left inset) and different experimental groups (upper inset).

(B) Pearson correlation heatmap of quantified protein intensities in all technical replicates (n=3) and biological replicates (n=3) from 36 LC-MS/MS runs for HPC. (#experimental group_technical replicate)

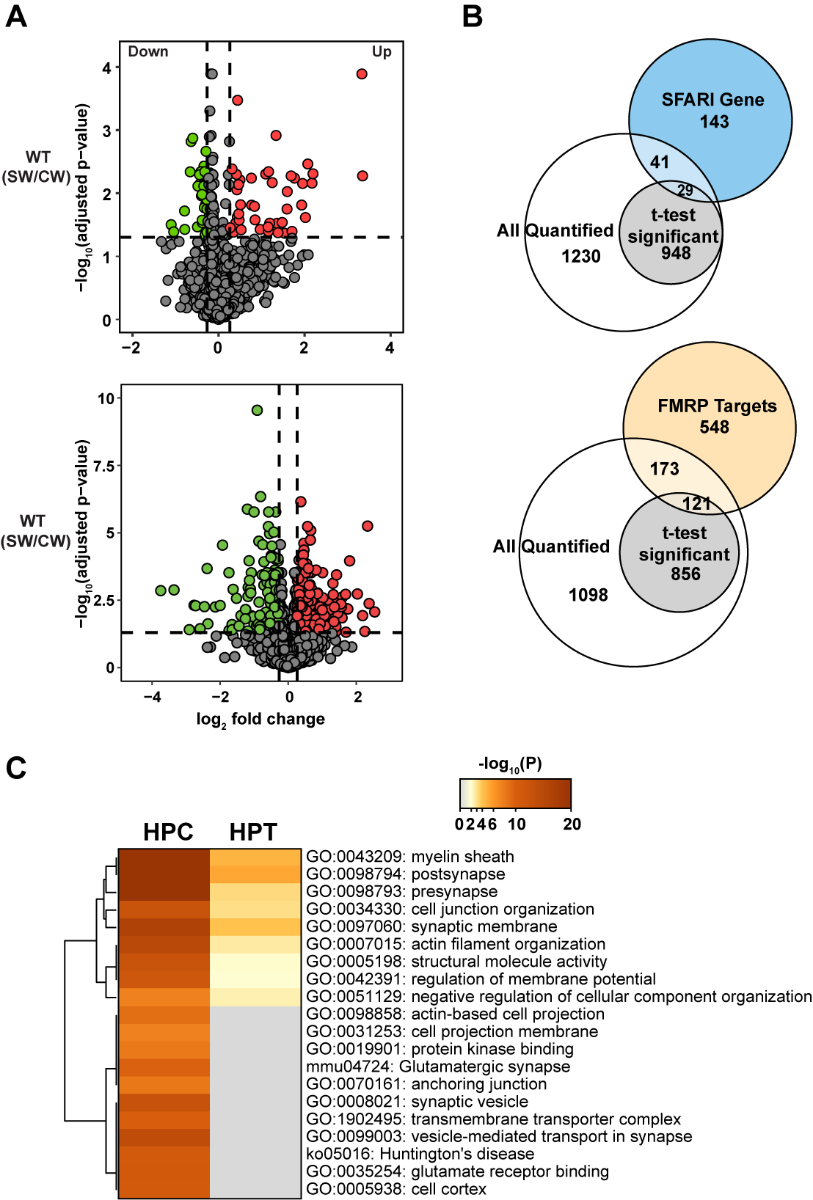

**Figure S6. Global Shotgun Proteomics Analysis of Mouse HPT and HPC.**

(A) Volcano plots showed protein expression level changes between soy protein versus casein protein diets in WT group for HPT (upper panel) and HPC (lower panel). Log_2_ protein fold changes are plotted against the negative log_10_ adjusted *p*-values. Points above the non-axial horizontal line represent significantly altered proteins (adjusted *p-*value < 0.05, Student’s *t*-test). Significantly down-regulated proteins are shown in green and up-regulated ones are shown in red (|protein fold change| > 1.2).

(B) Comparison of the genes from all quantified proteins and significantly changed proteins (adjusted *p-*value < 0.05, Student’s *t*-test) in HPC to gene entries from FMRP target dataset by Darnell et al (yellow), as well as from SFARI autism database (blue).

(C) Heatmaps generated using Metascape showing the significantly enriched (*p* value < 0.01) biological processes, cellular components, molecular functions and Kyoto encyclopedia of genes and genomes (KEGG) pathways for WT group in HPT and HPC. The most significant 20 terms are shown in the heatmap. Color coding indicates -log_10_ (*p* values). Rows are clustered based on their profile similarity.

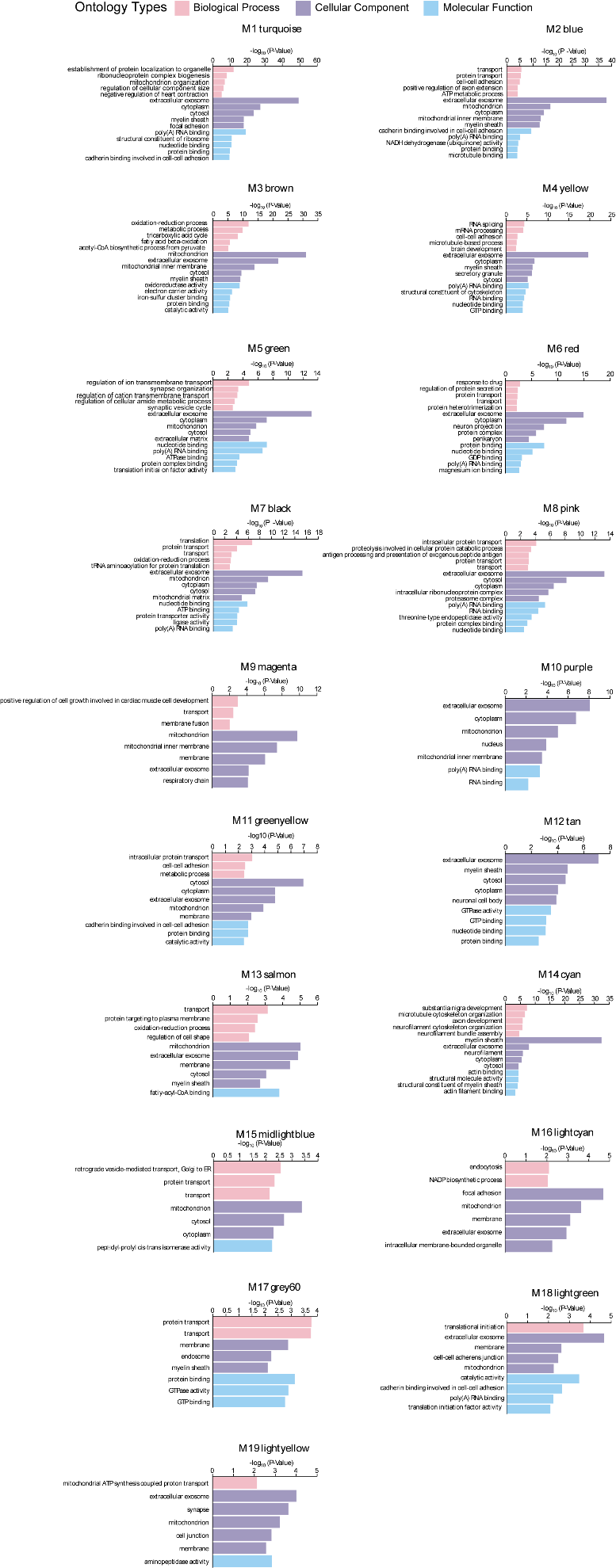

**Figure S7. GO analysis of HPT *Fmr1^KO^* group protein network modules.**

Gene ontology (GO) analysis was performed using DAVID bioinformatics resources to gain biological insight into each protein network module. Enrichment for a given ontology was shown as log_10_ *p*-values.

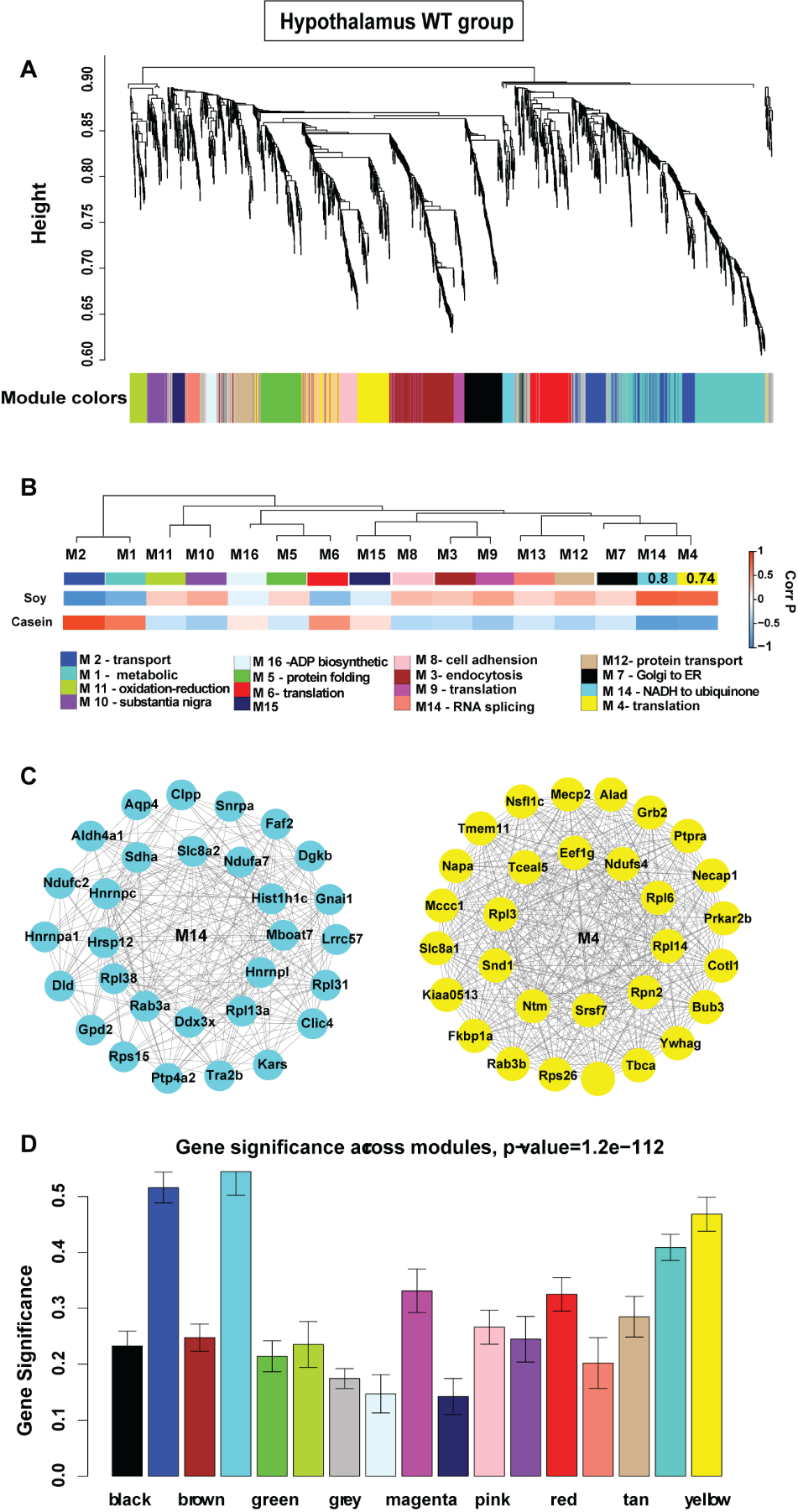

**Figure S8. WGCNA co-expression network analysis for HPT WT group proteins.**

(A) Protein clustering trees generated in WT group for HPT. Each ME is in non-gray.

(B) Protein correlation network consisting of 16 protein modules was generated from 1,866 proteins in WT group for HPT. Module eigenproteins, which represent the first PC of the protein expression within each module, were correlated with diet trait. Strength of positive (red) or negative (blue) correlation is shown by two-color heat map, with *P* values provided for all correlations with *P* < 0.05. Modules that showed a significant correlation with diet trait are labeled in correlation value. GO analysis of the proteins within each module clearly identified the biological functions, processes, and components associated with the module for most modules.

(C) Network of top 50 hub proteins in ME14, NADH to ubiquinone, (left) and ME4 translation (right) in WT group for HPT.

(D) Gene significance across modules for WT group proteins in HPT.

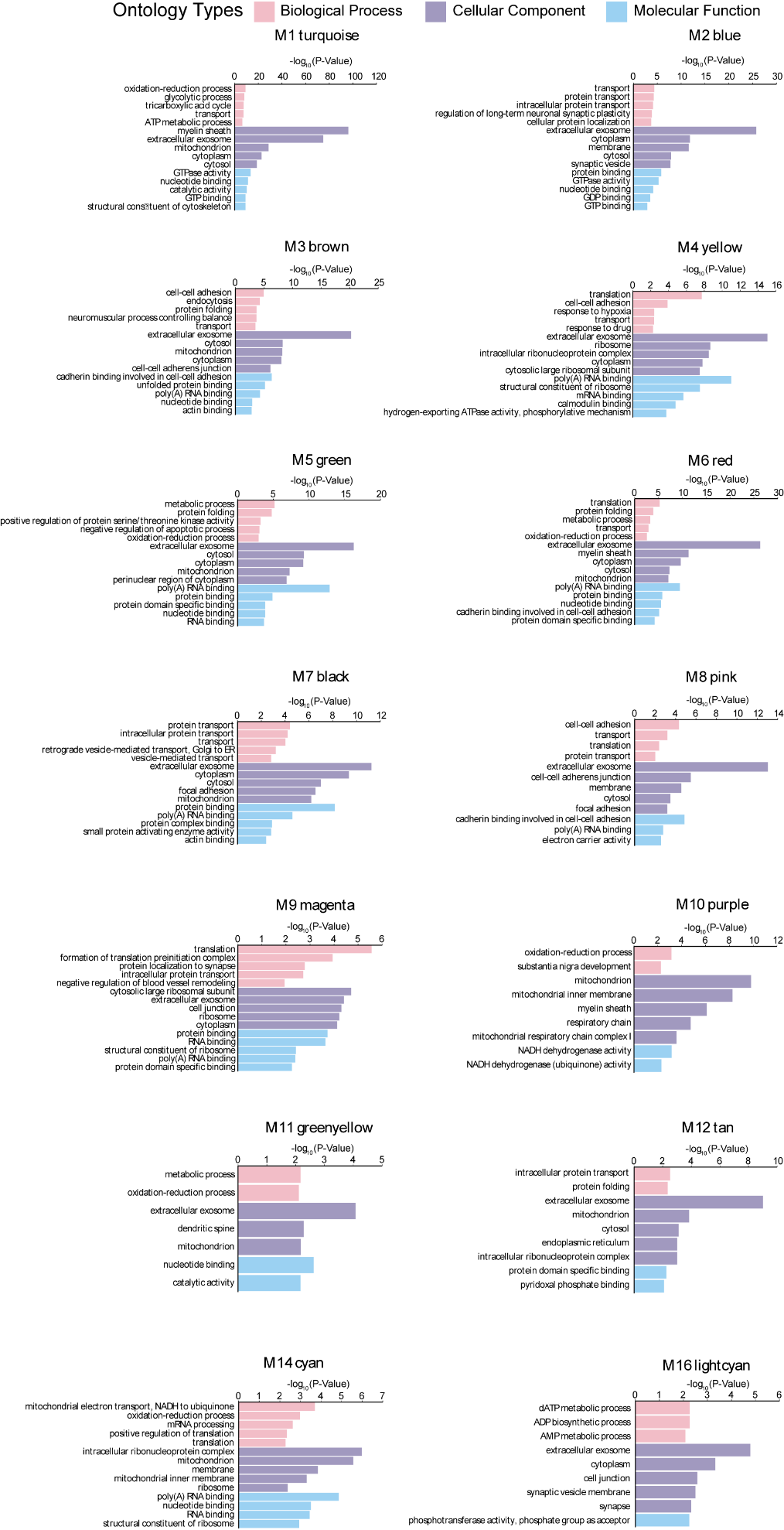

**Figure S9. GO analysis on HPT WT group protein network modules.**

Gene ontology (GO) analysis was performed using DAVID bioinformatics resources to gain biological insight into each protein network module. Enrichment for a given ontology was shown as log_10_ *p*-values.

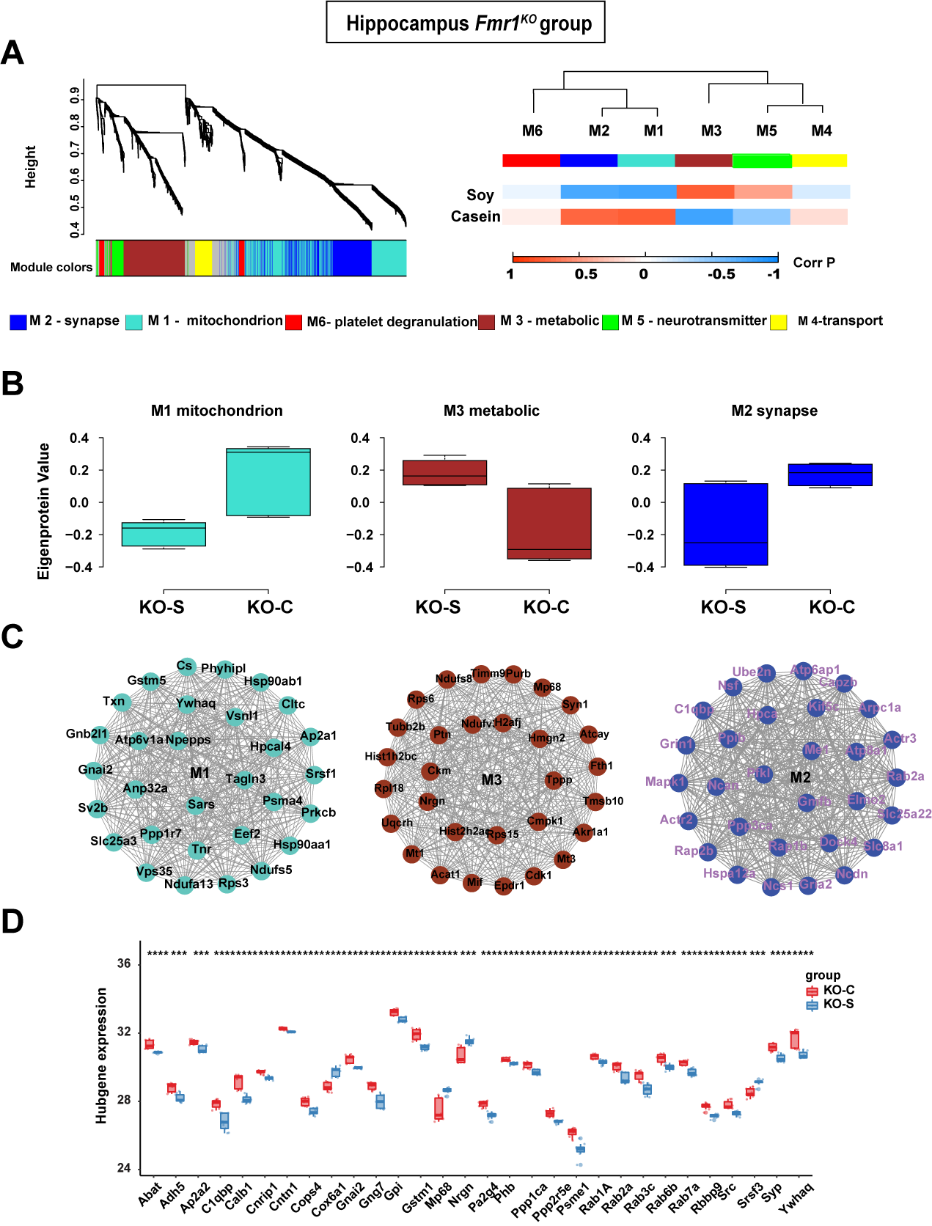

**Figure S10. WGCNA co-expression network analysis for HPC *Fmr1^KO^* group** **proteins.**

(A) Protein clustering trees generated in *Fmr1^KO^* group for HPC and each ME is in non-gray (left panel). Protein correlation network consisting of 6 protein modules was generated from 2,061 proteins in *Fmr1^KO^* group for HPC (right panel). Module eigenproteins, which represent the first PC of the protein expression within each module, were correlated with diet trait. Strength of positive (red) or negative (blue) correlation is shown by two-color heat map, with *P* values provided for all correlations with *P* < 0.05. Modules that showed a significant correlation with diet trait are labeled in correlation value. GO analysis of the proteins within each module clearly identified the biological functions, processes, and components associated with the module for most modules (bottom).

(B) ME level by casein- or soy-diet for ME1 mitochondrion (left), ME3 metabolic (middle) and ME2 synapse (right) that had significant correlation to two traits. Whiskers extend to data points that are less than 1.5* inter-quartile range away from 1^st^ and 3^rd^ quartile, respectively. The horizontal line shows the median.

(C) Network of top 50 hub proteins in ME1 mitochondrion (left), ME3 metabolic (middle) and ME2 synapse (right) in *Fmr1^KO^* group for HPC.

(D) Top 20 representative proteins selected in ME. Box plots represent the median, and 25th and 75th percentiles, and whiskers represent measurements to the 5th and 95th percentiles. Differences in protein levels were assessed by Wilcoxon-test (**p*-value<0.05, ***p*-value<0.001, ****p*-value<0.0001).

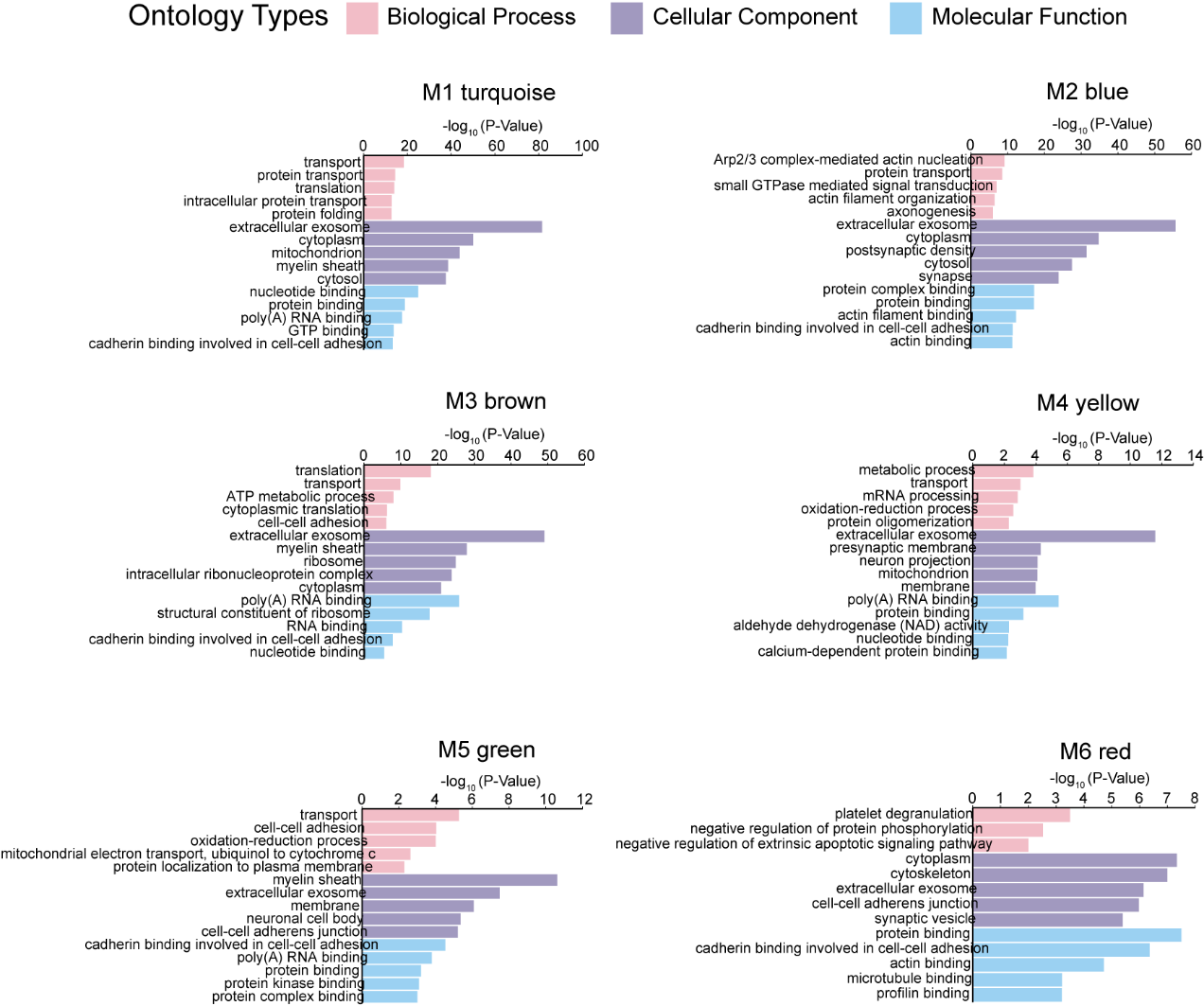

**Figure S11. GO analysis of HPC *Fmr1^KO^* group protein network modules.**

Gene ontology (GO) analysis was performed using DAVID bioinformatics resources to gain insight into the biological meaning of each protein network module. Enrichment for a given ontology was shown as log_10_ *p*-values.

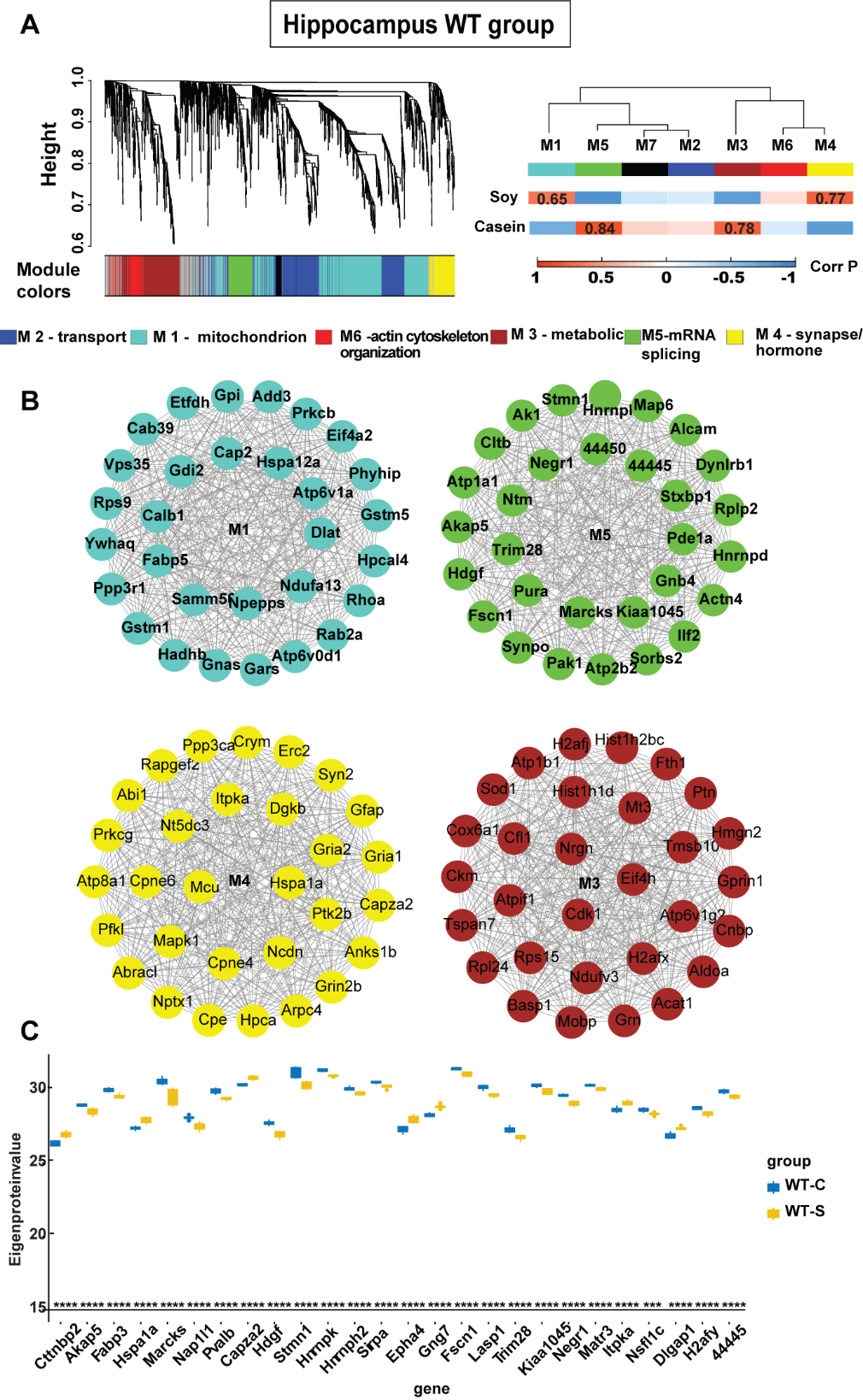

**Figure S12. WGCNA co-expression network analysis for HPC WT group proteins.**

(A) Protein clustering trees generated in WT group for HPC and each ME is in non-gray (left panel). Protein correlation network consisting of 7 protein modules was generated from 2,217 proteins in WT group for HPC (right panel). MEs were correlated with soy protein or casein protein diets. Strength of positive (red) or negative (blue) correlation is shown by two-color heat map, *P* values are provided for modules that showed a significant correlation with two traits. GO analysis of the proteins within each module clearly identified the biological functions, processes, and components associated with the module for most modules (bottom).

(B) Network of top 50 hub proteins in ME1 mitochondrion (upper left), ME5 mRNA splicing or processing (upper right), ME4 actin cytoskeleton organization (lower left) and ME3 metabolic (lower right) in WT group for HPC.

(C) Top 20 representative proteins selected in ME3. Box plots represent the median, and 25th and 75th percentiles, and whiskers represent measurements to the 5th and 95th percentiles. Differences in protein levels were assessed by Wilcoxon-test (**p*-value<0.05, ***p*-value<0.001, ****p*-value<0.0001).

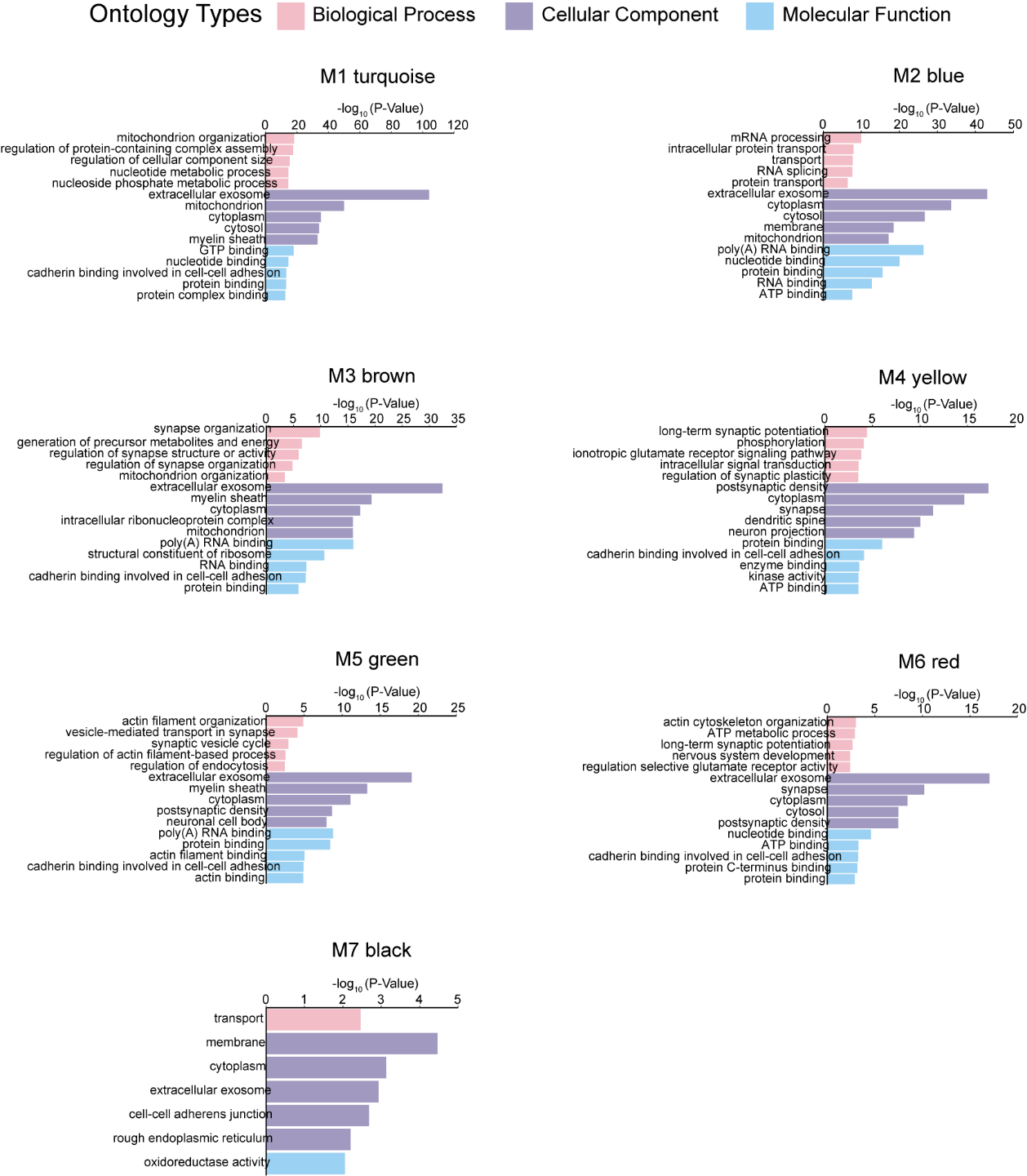

**Figure S13. GO analysis of HPC WT group protein network modules.**

Gene ontology (GO) analysis was performed using DAVID bioinformatics resources to gain insight into the biological meaning of each protein network module. Enrichment for a given ontology was shown as log_10_ *p*-values.

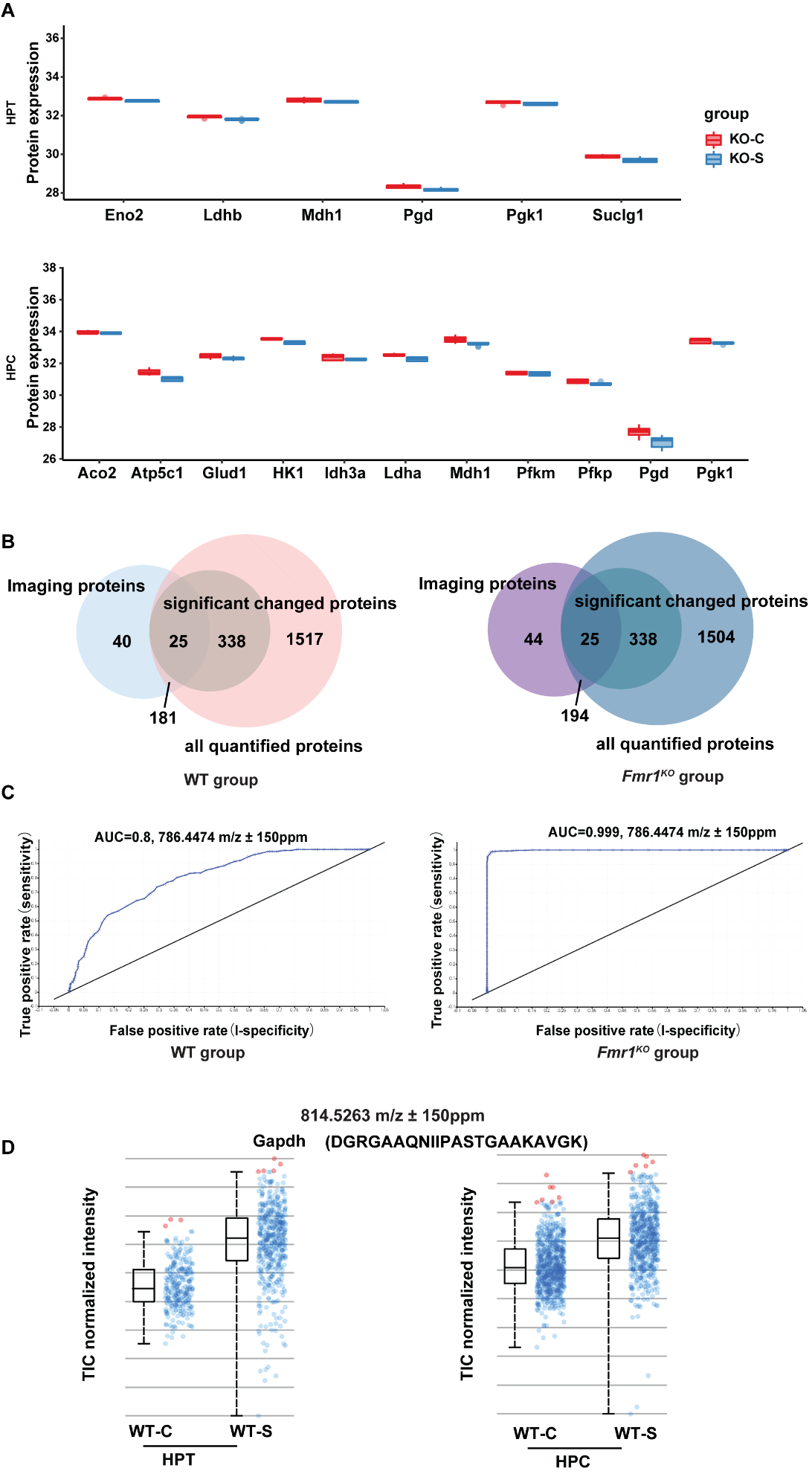

**Figure S14. Spatially resolved biomarkers compatible with shotgun proteomics analysis.**

(A) Enzymes involved in glycolytic and TCA cycle pathways are dysregulated by soy consumption in both HPT (upper panel) and HPC (lower panel) of FXS.

(B)Venn diagram showing the overlap among MS imaging annotated proteins, LC-MS/MS all quantified protein numbers, and significantly changed proteins in WT group (left panel) and *Fmr1^KO^* group (right panel) in HPT.

(C) Adjusted ROC analysis corresponding to Figure 7B in WT group (left panel) and *Fmr1^KO^* group (right panel) of HPT.

(D) Normalized intensity of total ion signals of the Gapdh protonated peptide in four cohorts in HPT (left panel) and HPC (right panel) of WT group. The boxplot points correspond to relative intensities of m/z 814.5263-related peaks taken from each spectrum in all images acquired.
